# Evolutionary and Developmental Specialization of Foveal Cell Types in the Marmoset

**DOI:** 10.1101/2023.12.10.570996

**Authors:** Lin Zhang, Martina Cavallini, Junqiang Wang, Ruiqi Xin, Qiangge Zhang, Guoping Feng, Joshua R. Sanes, Yi-Rong Peng

## Abstract

In primates, high-acuity vision is mediated by the fovea, a small specialized central region of the retina. The fovea, unique to the anthropoid lineage among mammals, undergoes notable neuronal morphological changes during postnatal maturation. However, the extent of cellular similarity across anthropoid foveas and the molecular underpinnings of foveal maturation remain unclear. Here, we used high throughput single cell RNA sequencing to profile retinal cells of the common marmoset (*Callithrix jacchus*), an early divergent in anthropoid evolution from humans, apes, and macaques. We generated atlases of the marmoset fovea and peripheral retina for both neonates and adults. Our comparative analysis revealed that marmosets share almost all its foveal types with both humans and macaques, highlighting a conserved cellular structure among primate foveas. Furthermore, by tracing the developmental trajectory of cell types in the foveal and peripheral retina, we found distinct maturation paths for each. In-depth analysis of gene expression differences demonstrated that cone photoreceptors and Müller glia, among others, show the greatest molecular divergence between these two regions. Utilizing single-cell ATAC-seq and gene-regulatory network inference, we uncovered distinct transcriptional regulations differentiating foveal cones from their peripheral counterparts. Further analysis of predicted ligand-receptor interactions suggested a potential role for Müller glia in supporting the maturation of foveal cones. Together, these results provide valuable insights into foveal development, structure, and evolution.

**Significance statement:** The sharpness of our eyesight hinges on a tiny retinal region known as the fovea. The fovea is pivotal for primate vision and is susceptible to diseases like age-related macular degeneration. We studied the fovea in the marmoset–a primate with ancient evolutionary ties. Our data illustrated the cellular and molecular composition of its fovea across different developmental ages. Our findings highlighted a profound cellular consistency among marmosets, humans, and macaques, emphasizing the value of marmosets in visual research and the study of visual diseases.

## Introduction

Diurnal (day-active) anthropoid primates, including monkeys, apes, and humans, are highly visual animals that perceive the world with high spatial and chromatic resolution (1). Nearly all of their high-acuity and most of their chromatic vision is mediated by a small, specialized central region of the retina called the fovea, which is absent from all other mammals and may be the only primate-specific structure in the mammalian brain (2–4) (Fig. 1*A*). Two observations dramatize the importance of the fovea. First, although the fovea occupies only ∼1% of the retinal surface, it supplies ∼50% of retinal input to the visual cortex (5, 6). Second, injuries or diseases that disable the fovea lead to devastating visual impairment whereas loss of far larger swaths of peripheral retina has much milder effects (7, 8).

**Figure 1.**
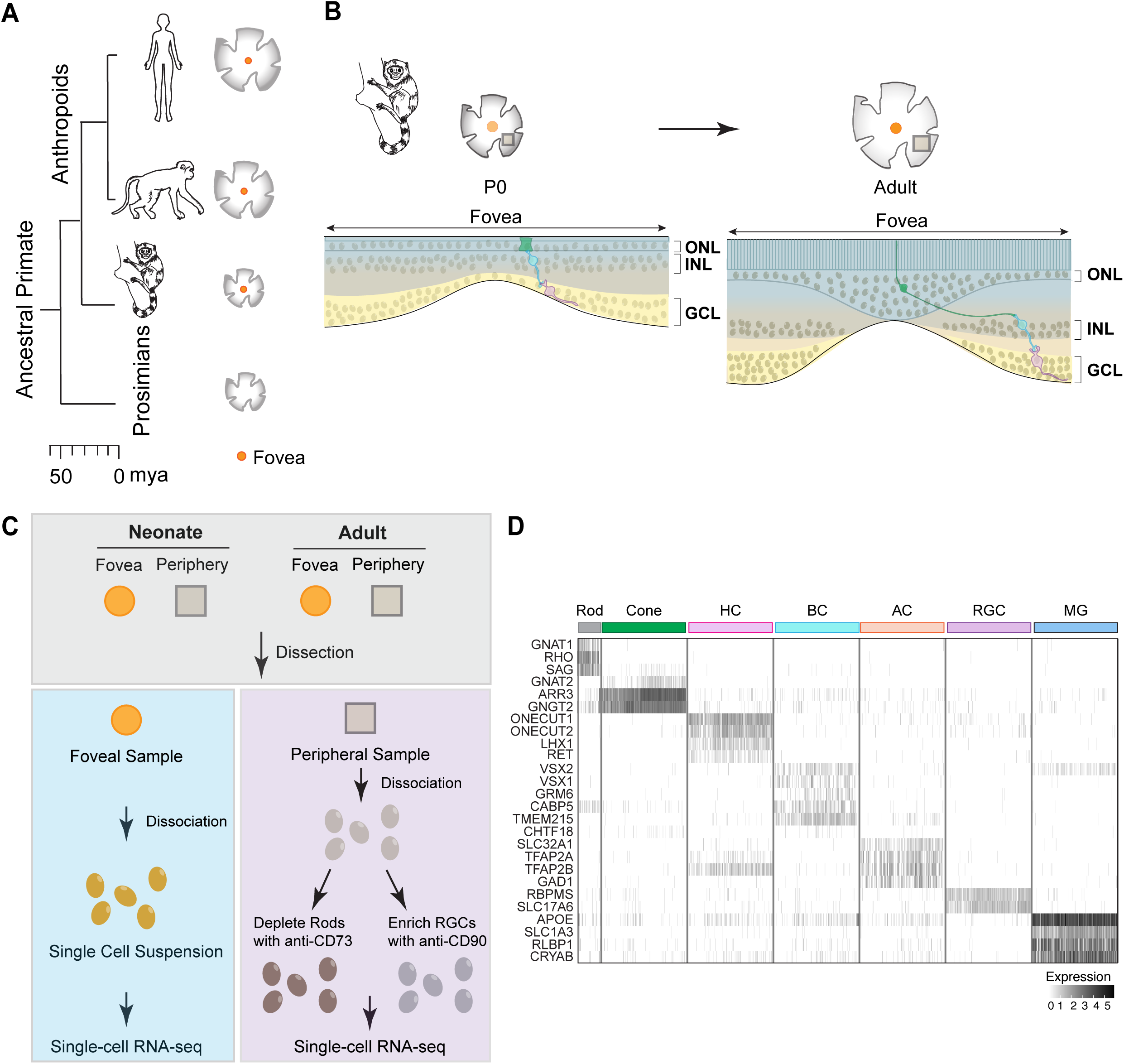
The evolutionary and developmental formation of the primate fovea. *(A)* Phylogenetic tree of selected primate species illustrating the evolutionary emergence of the fovea in anthropoids. The scale bar indicates the estimated divergence time in millions of years ago (MYA). *(B)* Sketches of the cellular arrangement in the marmoset fovea at the neonatal (left) and adult (right) stages. The midget pathway, consisting of a cone (green), a midget bipolar cell (blue), and a midget ganglion cell (purple) is demonstrated in both neonatal and adult fovea to highlight its developmental change. The morphological transformation of a foveal cone during postnatal maturation is particularly noteworthy. *(C)* Diagram summarizing the foveal and peripheral regions collected from neonatal and adult marmoset retinas, along with the experimental workflow for single-cell RNA-seq. Peripheral cells were depleted of rods using anti-CD73 antibody or enriched with RGCs using anti-CD90 antibody. *(D)* Heatmap displaying the expression of marker genes in individual cell classes in the neonatal marmoset fovea.

The fundamental cellular plan of the fovea is similar to that of the peripheral retina (9). In both cases, cells comprise 6 main classes, most of which can be divided into multiple types: photoreceptors (PRs), which sense light; interneurons (horizontal, bipolar, and amacrine cells), which process visual information relayed from PRs; retinal ganglion cells (RGCs), which integrate input from interneurons and send axons through the optic nerve to the rest of the brain; and a single endogenous glial type called the Müller glia (9, 10). However, the fovea has modified this basic plan with a variety of structural and functional specializations: (1) The fovea, and particularly its central portion, the foveola, are depressed, forming a shallow pit (5, 6) (Fig. 1*B*). (2) The inner nuclear and ganglion cell layers, containing interneurons and RGCs, are displaced, presumably to reduce light scattering. (3) Nearly all PRs in the foveal center are cones whereas >90% of PRs in the peripheral retina are rods (11, 12). (4) Consistent with cone dominance, bipolar cells that receive input primarily from rods (rod bipolar cells) are sparse in the fovea but the major bipolar type in the periphery (5, 13). (5) The outer segments of foveal cones, which contain the visual pigment (opsin) are far longer and thinner than those in peripheral retina (14). The length enhances sensitivity while the small diameter improves spatial resolution. (6) Likewise, axons of foveal cones are longer than those of peripheral cones, enabling them to reach displaced bipolar and horizontal cell dendrites in the inner nuclear layer (15). (7) Excitatory drive to most foveal RGCs arises from a single PR, whereas peripheral RGCs may receive input from dozens of PRs (5, 6, 11). This arrangement maximizes the spatial resolution of foveal RGCs, albeit at the expense of sensitivity.

Although the anatomical and physiological bases of these specializations have been documented in detail, their molecular correlates remain largely unknown. Here, we have used high throughput single cell transcriptome profiling (scRNA-seq) to address issues relating to the cell types, development, and evolution of the fovea in the common marmoset (*Callithrix jacchus*). The marmoset is well-suited for this inquiry for three reasons. First, its small size, relatively short gestation time and genetic accessibility make it useful for developmental studies (16, 17). Second, these features have enabled numerous recent studies of the marmoset visual system, so much is now known about its retina (18–23). Third, New World (Platyrrhine) primates including marmosets diverged early in anthropoid evolution (35-40 million years ago) from Old World (Catarrhine) primates, such as macaques, apes, and humans (16, 24, 25) (Fig. 1*A*). Thus, comparison of marmoset fovea with those of macaques and humans can provide insights into how it arose in primates.

Our aim in the work reported here was to learn more about how the fovea arises both in phylogeny, as foveated primates arose from their non-foveated ancestors, and during ontogeny, as the fovea differentiates from an initially uniform retina. To this end, we generated cell atlases from neonatal and adult marmoset fovea and peripheral retina and compared them to each other and to atlases we had previously generated from adult human and macaque fovea and peripheral retina (26, 27). We found that over 90% of cell types were shared across regions, ages, and species. However, in each case we found large numbers of genes differentially expressed by shared types. We used a variety of computational methods to gain insight into the biological significance of these differences. Our results highlighted foveal cone photoreceptors and Müller glia as cell types that differ markedly between neonates and adults and between periphery and fovea. Further analysis of these differences suggested that factors derived from Müller glia might promote the maturation of foveal cones.

## Results

### Cell atlas of the adult marmoset retina

We generated a cell atlas of the adult (>2-year-old) marmoset fovea and peripheral retina. Cells were dissociated for high throughput scRNA-seq using the 10X Genomics platform (28) (Fig. 1*C*). Foveal cells were profiled without pretreatment. Peripheral cells were treated with anti-CD73 to deplete rod PRs or with anti-CD90 to enrich RGCs, using methods we had developed in studies of macaque (26). Despite the use of distinct treatments for foveal and peripheral samples, the molecular complexities of cells in both regions are comparable (*SI Appendix*, Fig. S1*A*). We also performed single nucleus RNA-seq (snRNA-seq) on the peripheral retina. After filtering out poor quality cells or nuclei, we obtained 29,169 foveal and 15,098 peripheral high-quality transcriptomes (*SI Appendix*, Table S1).

Based on unsupervised clustering methods, we divided the foveal cells into six cell classes, which we annotated using known class-specific marker genes (26, 27, 29, 30): PRs, horizontal cells (HCs), bipolar cells (BCs), amacrine cells (ACs), RGCs, and non-neuronal cells (NN) (Fig. 1*D*). From each cell class, we further clustered individual cell types. In total, we identified a total of 68 types: 3 PRs, 2 HCs, 13 BCs, 30 ACs, 16 RGCs and 4 NN types (Fig. 2*A*, *SI Appendix*, Table S2). Biases associated with specific animal or sample batches were negligible after batch correction (*SI Appendix*, Fig. S1*B*).

**Figure 2.**
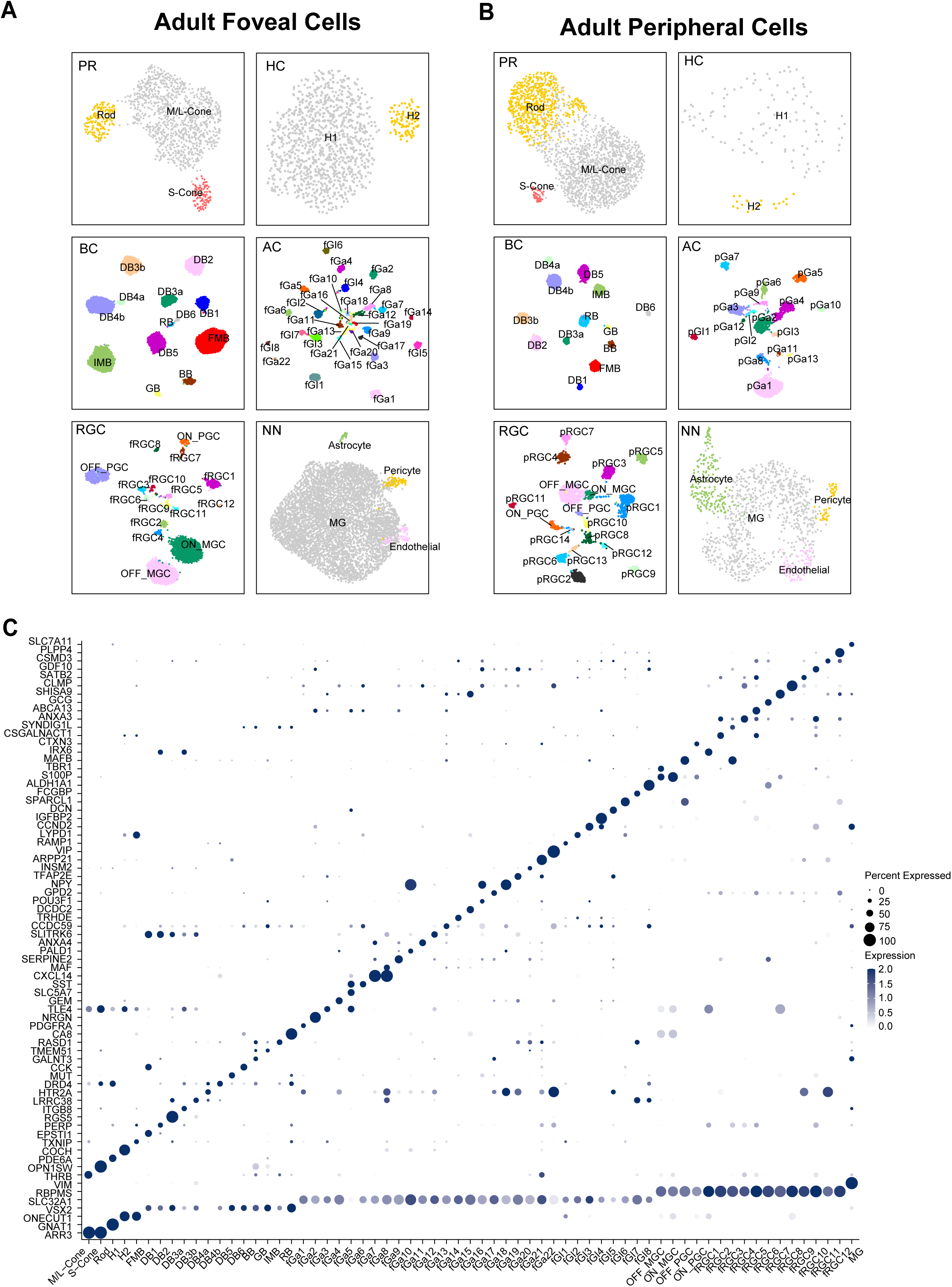
Cell atlas of the adult marmoset retina. *(A)* Uniform Manifold Approximation and Projection (UMAP) visualization of cell types from individual cell classes (PR, photoreceptors; HC, horizontal cells; BC, bipolar cells; AC, amacrine cells; RGC, retinal ganglion cells; NN, non-neuronal cells) in the fovea of adult marmoset. *(B)* UMAP visualization of cell types from individual cell classes in the peripheral retina of adult marmoset. *(C)* Dot plot showing the expression of marker genes for individual foveal cell classes (bottom seven rows) and types (remaining rows).

We verified the clustering result in two ways. First, we asked to what extent all these cell types can still be identified without prior separation into cell classes. To this end, we analyzed all foveal cells together and clustered using an iterative unsupervised method with a maximum resolution (See Materials and Methods). Cell types identified by this method corresponded to those described above (*SI Appendix*, Fig. S1*C*). Second, we analyzed differentially expressed genes and identified markers that are specifically expressed by individual cell types (Fig. 2*C*, *SI Appendix*, Fig. S2). Thus, multiple approaches identified a set of 68 transcriptomically distinguishable foveal cell types.

We clustered peripheral retinal cells obtained by scRNA-seq using similar methods but supplemented them with RGCs and HCs from the snRNA-seq dataset as these two classes had fewer numbers of cells than others (*SI Appendix*, Table S1). We integrated scRNA-seq and snRNA-seq data by canonical correlation analysis (CCA) to overcome potential expression-level shift caused by different preparation methods (31) (*SI Appendix*, Fig. S1*D* and *E*). We identified a total of 56 types including 3 PRs, 2 HCs, 13 BCs, 16 ACs, 18 RGCs, and 4 NN types (Fig. 2*B*). The smaller number of AC types from periphery compared to fovea likely reflects the poor recovery of glycinergic ACs with CD90 selection (26). Like the foveal dataset, we were able to classify all the cell types using high resolution and identified specific marker genes for individual cell types (*SI Appendix*, Fig. S1*E* and S3). All peripheral types were shared with foveal types in the adult retina (see below).

## Evolutionary modification of foveal cell types across anthropoids

Marmosets diverged from a common ancestor prior to macaques and humans, which are more closely related to each other than to marmosets (16, 25) (Fig. 1*A*). We compared foveal cell types of the three species as one way of asking whether all foveal cell types were present in the primate ancestor. We performed three comparisons. First, we pooled a maximum of 200 cells per type from each species and used CCA to integrate a total of 23,966 cells, followed by unsupervised clustering. All cells fell into the six canonical cell classes, which can be further classified into known subclasses (e.g., GABAergic and Glycinergic ACs) (32) or types (*SI Appendix*, Fig. S4*A*). For RGCs, we distinguished midget, parasol, and intrinsically photosensitive (ip) RGCs (18, 33, 34), but group all other minor types as a single subclass. These 26 cell types/subclasses contain cells from all three species without species bias (Fig. 3*A*, *SI Appendix*, Fig. S4*B* and *C*). We found both conserved and species-specific marker genes in homologous types and subclasses (Fig. 3*A*, *SI Appendix*, Fig. S4*D*).

**Figure 3.**
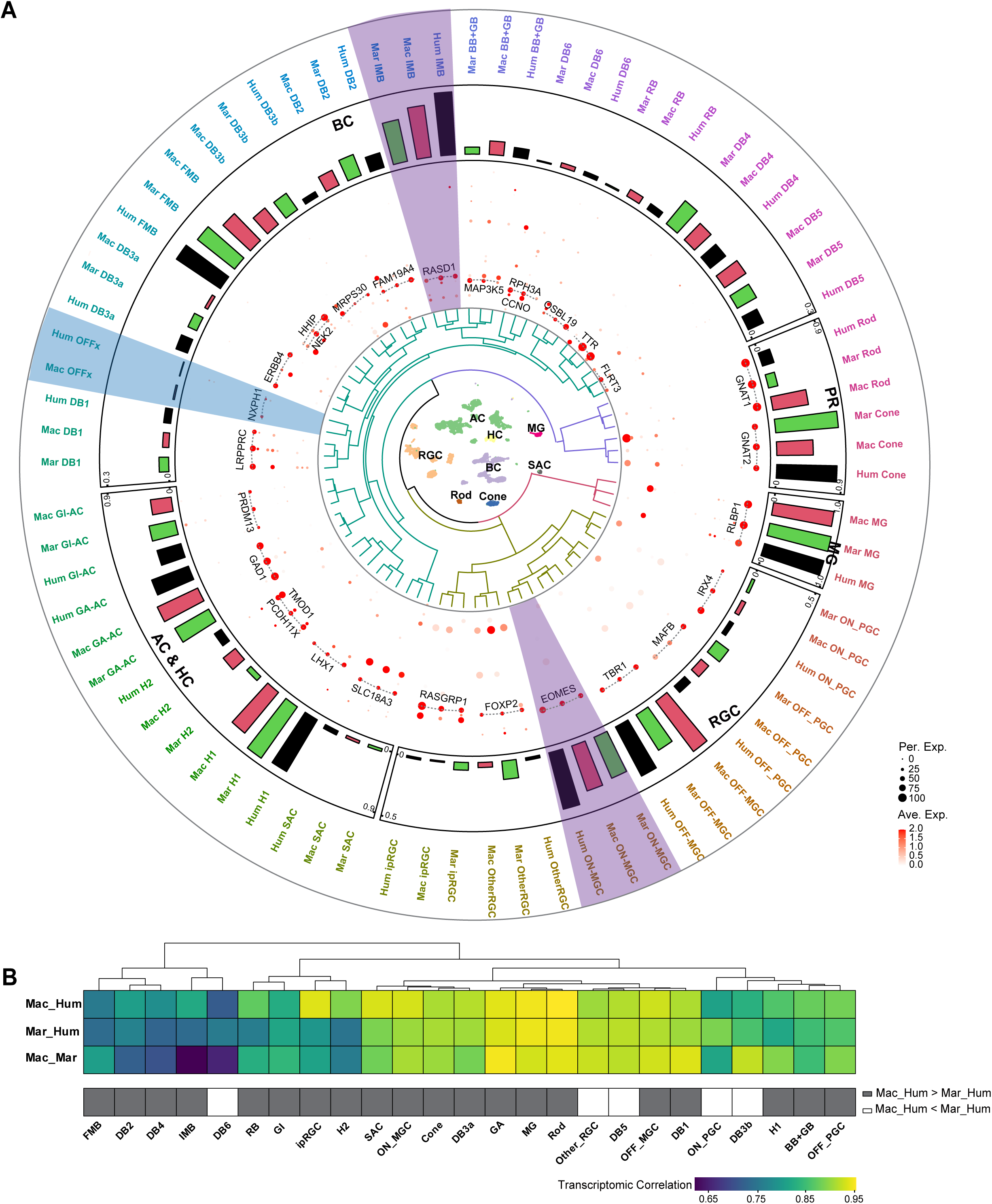
Conserved and divergent features of foveal cell types across three anthropoid primate species. *(A)* Overview of foveal cell types via comparative analysis of scRNA-seq data from human, macaque, and marmoset. Depicting the plot from inner-to outermost as follows: UMAP of homologous cell types integrated from 23,966 cells across three species; dendrogram of hierarchically related homologous cell types; dot plot showing the expression of conserved marker genes among homologous cell types across species; bar plot showing proportions of cell types within individual cell classes across species, with the maximum proportion within each class indicated at the top edge of the bar plots; 77 homologous cell types. Purple bars highlight ON MGC and IMB, and blue bar highlights the OFFx BC. *(B)* Transcriptomic correlation of homologous cell types between each pair of the three species via Pearson correlation. Grey-filled boxes indicate cell types with a higher correlation between macaque and human, as opposed to marmoset and human, while open boxes indicate the opposite pattern.

Second, we compared the transcriptomic distance among homologous cell types across the three primates. Hierarchical clustering of homologous cell types showed that most human and macaque types are more similar to each other than either is to the marmoset type (Fig. 3*A*). We also applied a transcriptomic mapping method to match types based on highly variable genes (HVGs) shared between species. Nearly twice as many marmoset cells favorably matched to macaque as to human types in BC, RGC, and AC classes (*SI Appendix*, Fig. S4*E*).

Third, we analyzed the transcriptomic convergence of homologous foveal cell types based on the correlated expression of HVGs (Fig. 3*B*). All pair-wise comparisons showed >61% correlations, with correlation of homologous types between macaques and humans higher than those in marmoset-human comparisons in 20/25 cases (Fig. 3*B*). Thus, the transcriptomic relationship among foveal cell types mirrors the evolutionary distance among primates (Fig. 1*A*) (25).

We also compared abundance of foveal cell types among the three species (Fig. 3*A*, *SI Appendix*, Table S3). The abundance of most types was similar across species, but several species-specific enrichments emerged. (1) The abundance of two major cell types in the ON midget pathway–invaginated midget bipolar (IMB) and ON midget ganglion cells (ON_MGC) varies in the order: human > macaque > marmoset, suggesting enrichment of the ON midget pathway in the fovea. (2) The fraction of cones among all photoreceptors is higher in the human and marmoset fovea than in the macaque fovea. (3) Although the OFFx BC type is absent in marmosets, it has been identified morphologically in rodents and transcriptomically in several other species, including macaques and humans (26, 35–37). This pattern suggests that the BC1b/OFFx type has been lost in marmosets. Altogether, these results demonstrated that primate foveas share a diverse pool of conserved cell types, but that evolutionary modifications are present in the cellular and molecular composition of foveal cell types.

### Cell atlas of the neonatal marmoset retina

Next, we generated cell atlases from neonatal marmoset fovea and peripheral retina; similar to the adult retina, we profiled all cells from fovea but depleted rods or enriched RGCs from peripheral samples. We did not observe expression-level differences caused by different preparation methods (*SI Appendix*, Fig. S5*A*, *B*, and *D)*. From 21,675 high-quality single cell foveal transcriptomes, we identified 65 cell types: 3 PRs, 2 HCs, 13 BCs, 28 ACs, 15 RGCs, and 4 NN types; from 22,846 high-quality peripheral transcriptomes, we identified 68 types: 3 PRs, 2 HCs, 13 BCs, 31 ACs, 16 RGCs, and 3 NN types (Fig. 4*A* and *B*, *SI Appendix*, Table S1). Similar to the adult cell atlas, all cell types identified within individual classes could also be identified through a global clustering of all the cells (*SI Appendix*, Fig. S5*C* and *E*), and we were able to identify specific marker genes for each cell type (Fig. 4*C*, *SI Appendix*, Fig. S6 and S7).

**Figure 4.**
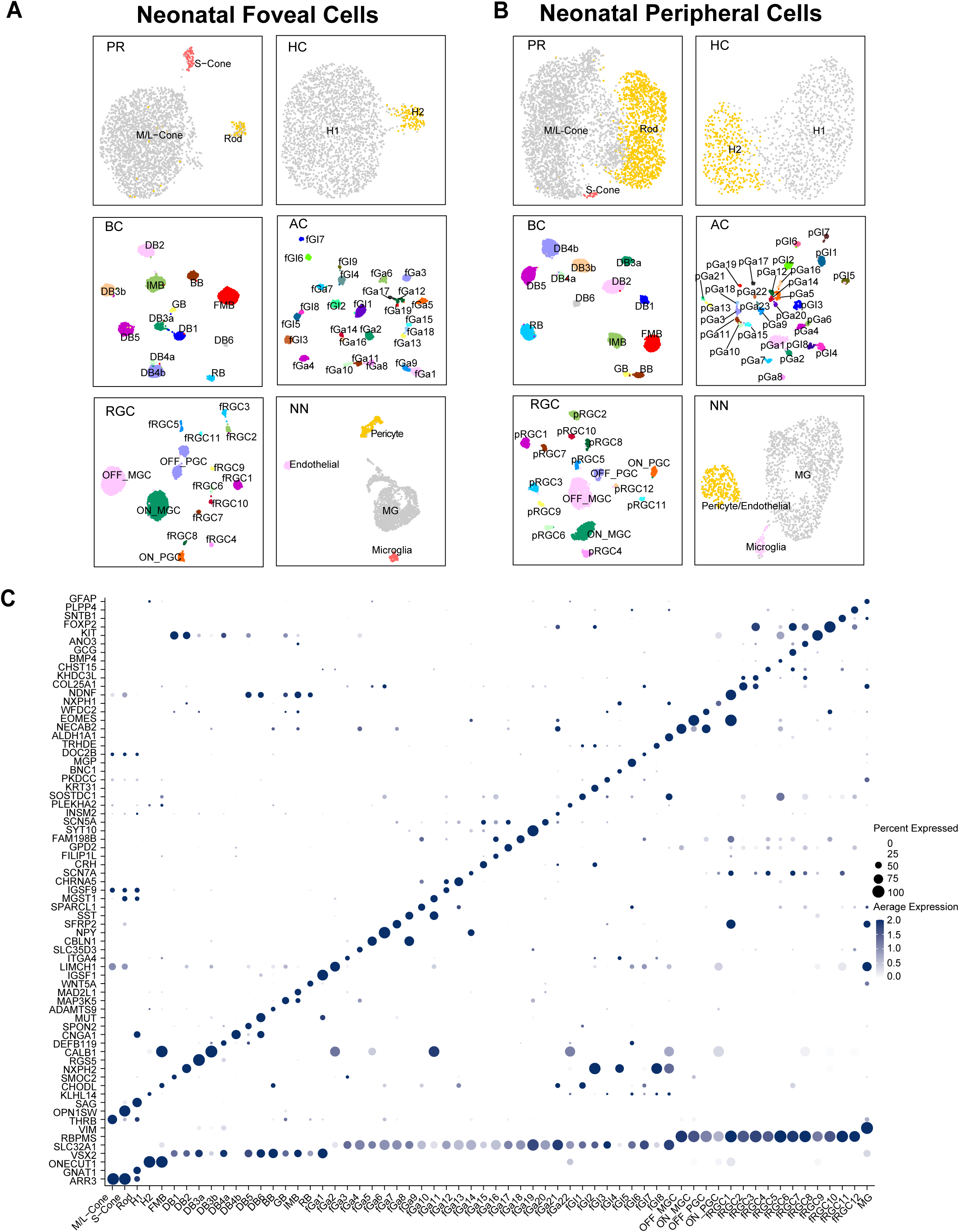
Cell atlas of the neonatal marmoset retina. *(A)* UMAP visualization of cell types from individual cell classes (PR, photoreceptors; HC, horizontal cells; BC, bipolar cells; AC, amacrine cells; RGC, retinal ganglion cells; NN, non-neuronal cells) in the fovea of neonatal marmoset. *(B)* UMAP visualization of cell types from individual cell classes in the peripheral retina of neonatal marmoset. *(C)* Dot plot showing the expression of marker genes for individual foveal cell classes (bottom seven rows) and types (remaining rows).

**Figure 5.**
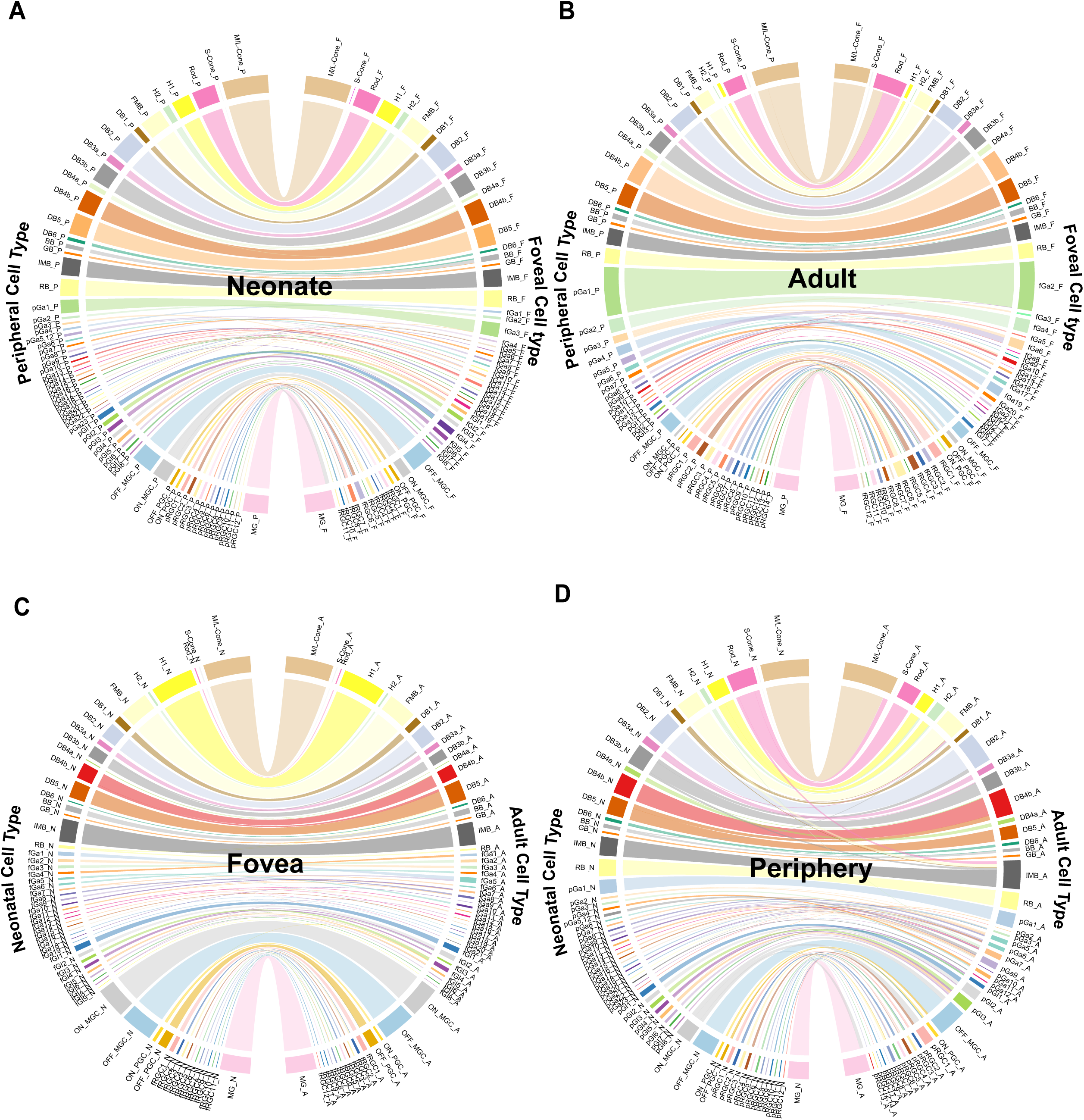
Transcriptomic correspondences of cell types across retinal regions and across developmental stages. *(A)* Chord diagram showing transcriptomic correspondence between foveal and peripheral cells in the neonatal stage. Each line maps a peripheral cell type (with suffix “_P”) to the corresponding foveal type (with suffix “_F”), and the width of each bar reflects the cell number. *(B)* Chord diagram showing transcriptomic correspondence between foveal and peripheral cells in the adult stages. Each line maps a peripheral cell type (with suffix “_P”) to the corresponding foveal type (with suffix “_F”), and the width of each bar reflects the cell number. *(C)* Chord diagram showing transcriptomic correspondence between neonatal and adult retinal cells in the fovea. Each line maps a neonatal cell type (with suffix “_N”) to the corresponding adult type (with suffix “_A”), and the width of each bar reflects the cell number. *(D)* Chord diagram showing transcriptomic correspondence between neonatal and adult retinal cells in the peripheral retina. Each line maps a neonatal cell type (with suffix “_N”) to the corresponding adult type (with suffix “_A”), and the width of each bar reflects the cell number. Cell types are ordered from top to bottom as PRs, HCs, BCs, ACs, RGCs and MG. The detailed correspondence is listed in *SI Appendix*, Table S4. The names of neonatal and adult cell types are consistent with those identified in Fig. 2 and 4.

### Cell types are shared between the fovea and peripheral retina and fully specified by birth

We next compared cell types across regions and ages. We applied a transcriptomic mapping method based on a multi-class classification framework using HVGs shared between pairs of datasets (26, 38) (see *SI Appendix*, Materials and Methods). For this analysis, we included Müller glia (MG) but excluded other NN types. There was a nearly complete correspondence between foveal and peripheral types in both neonates and adults (Fig. 5*A* and *B*). However, the low recovery of AC types with the CD90 enrichment method in the adult peripheral sample led to a one-to-multiple match in several cases (Fig. 5*B*, *SI Appendix*, Fig. S5D, Table S4). We next asked whether cell types are specified in neonates. Using the same transcriptomic mapping methods, we found that all 62 foveal cell types in the neonatal retina have corresponding types in the adult fovea. Out of these, 59/62 types had a 1:1 match, while two AC types and one RGC type showed a 2:1 match, suggesting a developmental maturation to further diversify these types (Fig. 5*C*, *SI Appendix*, Table S4). Similarly, all 66 peripheral cell types in the neonatal retina correspond to adult peripheral cell types, despite a multiple-to-one match observed in several cases due to the insufficient sampling of adult peripheral AC types (Fig. 5*C* and *D*, *SI Appendix*, Table S4). Thus, the fovea and peripheral retina share most if not all cell types, and nearly all cell types are present and molecularly specified by birth.

### Distinct developmental paths of cell types in the fovea and peripheral retina

Knowing that most cell types are present across regions and ages, we next analyzed regional (foveal versus peripheral region) and developmental (neonatal versus adult stage) differences between their transcriptomes. To assess the regional differences, we first integrated the cell types shared between the two regions at each age (Fig. 6*A* and *C*, *SI Appendix*, Tables S5). We then used two methods to calculate cell type–specific regional scores. The first was based on a signed gene-set enrichment analysis (sGSEA), which calculates a normalized enrichment score (NES) for genes differentially expressed between corresponding cell types (*SI Appendix*, Fig. S8*A* and *B*) (see *SI Appendix*, *Materials and Methods*). A higher NES indicates a greater regional difference. At both ages, only 3-4 cell types, such as MG and M/L-cones, showed significantly high NES values (Fig. 6*A* and *B*). Second, we calculated the earth mover’s distance (EMD) (39) scores to quantify the similarity between the density distributions of foveal and peripheral cells in terms of the expression of differential expression genes (*SI Appendix*, Fig. S8 *E* and *F*). A higher EMD score indicates a greater transcriptomic divergence between the corresponding foveal and peripheral type. Consistent with the sGSEA method, only 24% of neonatal cell types and 26% of adult cell types showed a regional difference above the mean change (Fig. 6 *B* and *D*). Therefore, using both methods, we found that the regional difference was not universally present in all cell types, but rather more significant in a small subset of cell types compared to others. Furthermore, we compared the regional differences of all cell types across ages by ranking their statistical significances, transforming the NES into a False Discovery Rate (FDR)-adjusted P value (see *SI Appendix*, Materials and Methods). We found that in adults, most cell types exhibited more significant regional differences than in the neonates, as determined by -log10FDR values (*SI Appendix*, Fig. S8*C*). Thus, transcriptomic differences between corresponding foveal and peripheral cell types increase during development.

**Figure 6.**
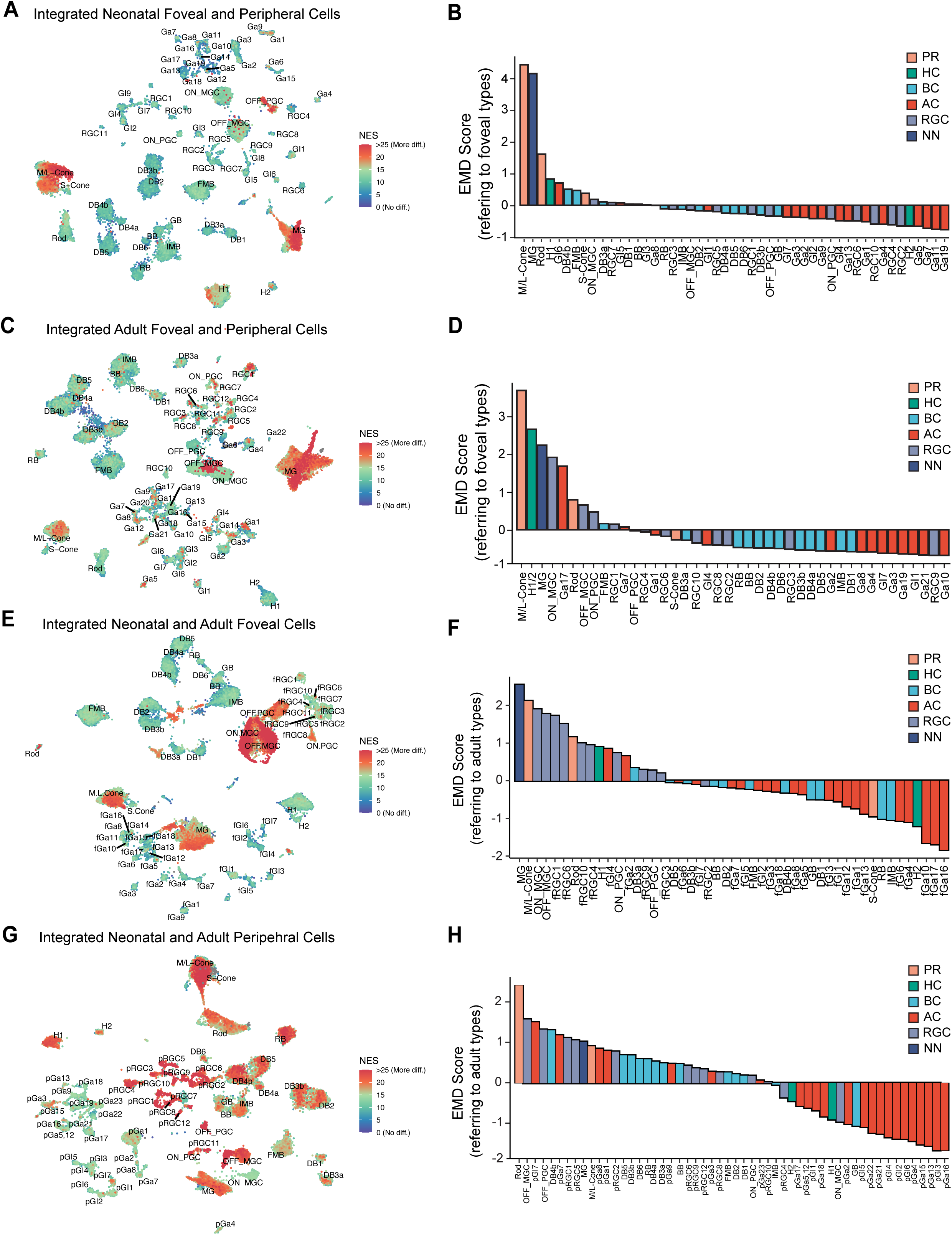
Transcriptomic comparisons of cell types across retinal regions and across developmental stages. *(A)* UMAP visualization of regional differences measured by normalized enrichment scores (NES) in individual foveal and peripheral cells, integrated at the neonatal stage. *(B)* Bar plot showing regional differences measured by scaled Earth Mover’s Distance (EMD) scores in integrated foveal and peripheral cell types at the neonatal stage. *(C)* UMAP visualization of regional differences measured by NES scores in individual foveal and peripheral cells, integrated at the adult stage. *(D)* Bar plot showing regional differences measured by EMD scores in integrated foveal and peripheral cell types at the adult stage. *(E)* UMAP visualization of developmental changes measured by NES scores in individual neonatal and adult cells, integrated at the foveal region. *(F)* Bar plot showing developmental changes measured by EMD scores in integrated neonatal and adult cell types at the foveal region. *(G)* UMAP visualization of developmental changes measured by NES scores in individual neonatal and adult cells, integrated at the peripheral region. *(H)* Bar plot showing developmental changes measured by EMD scores in integrated neonatal and adult cell types at the peripheral region. The NES values are divided into three ranges: Low (0–10), Medium (10–20), High (above 20). The correspondence between the names of integrated cell types and those at each stage are listed in *SI Appendix*, Table S5 for A. and C., and in *SI Appendix*, Table S6 for E. and G.

We then assessed the developmental differences between the fovea and peripheral retina. We separately integrated foveal and peripheral cell types across ages and again used sGSEA and EMD as measures (*SI Appendix*, Table S6). In the fovea, only a few cell types exhibited high developmental scores. In contrast, a large proportion of peripheral cell types showed high developmental scores (Fig. 6*E* and *G*). Similarly, using the EMD calculation, over 52% of cell types in the peripheral retina showed differences above the mean change, while only 26% of cell types in the fovea underwent significant developmental changes (Fig. 6*F* and *H*). Moreover, although most foveal cell types show a lesser significance in developmental change, as judged by the -log10FDR rankings, certain foveal cell types stood out with comparable significance comparable to the top peripheral peers (*SI Appendix*, Fig. S8*D*). These results demonstrate that there is a global maturation of cell types in the peripheral retina, indicating the peripheral retina is generally immature at birth; however, select foveal cell types still undergo substantial maturation processes during postnatal development.

### Conserved and divergent regional differences in cones and Müller glia across primates

All four analyses – sGSEA and EMD by region and by age, highlighted cones and MG – they showed the highest degree of regional difference at both ages and were among the top changed cell types across ages in the fovea (Fig. 6, *SI Appendix*, Fig. S8*C* and *D*). We therefore focused on these two cell types for our next analyses. Our initial investigation assessed the extent to which regional differences in cones and Müller glia are shared with those in humans and macaques. We curated a set of differentially expressed genes (DEGs) in these two cell types, comparing the fovea to the peripheral retina across three primates. We detected many DEGs that are shared among the three species, suggesting conserved roles for these genes in the regional specialization (*SI Appendix*, Fig. S9*A*-*D*). However, each species displayed a larger number of unique DEGs, indicative of molecular divergence and adaptation specific to their foveal function (*SI Appendix*, Fig. S9*A and C*).

### Biological pathways during the development of foveal cones

A hallmark of postnatal maturation in foveal cones is their drastic morphological changes they undergo. At birth, foveal cones are immature cuboidal cells arranged in one or two layers without obvious outer segments or axons (40, 41). As the fovea matures, they migrate centripetally, pack into ten somatic layers, develop long, slender outer segments and extend long axons (42) (Fig. 1*B*). To investigate the molecular changes associated with these development alterations, we pooled all 7,198 cones (foveal and peripheral from neonates and adults) and performed an unsupervised clustering of the dataset. They formed four discrete clusters, precisely divided by age and region (Fig. 7*A*).

**Figure 7.**
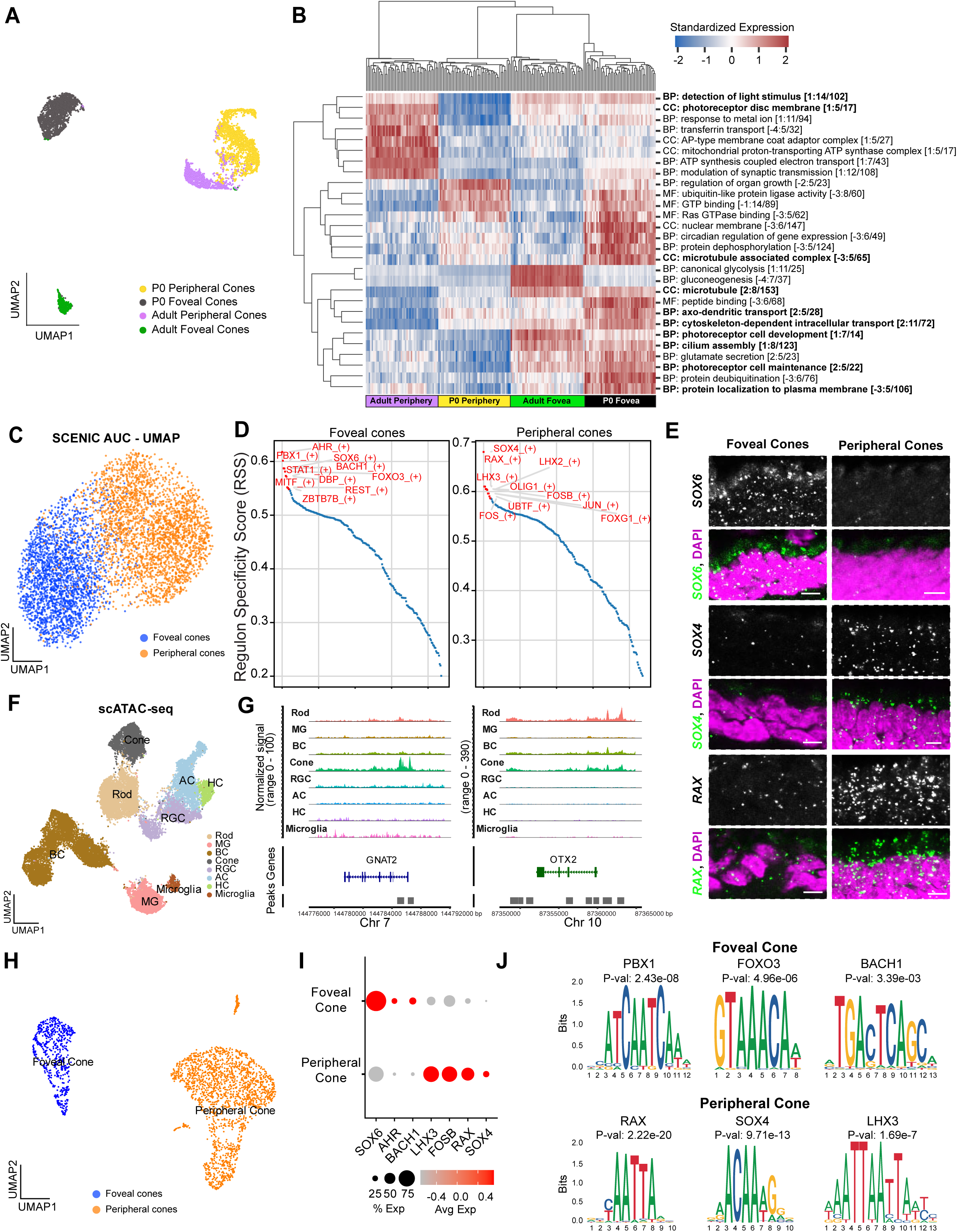
Distinct biological pathways and transcriptional landscapes associated with distinct cone types. *(A)* UMAP visualization of four cone clusters, each representing either foveal or peripheral cones at either neonatal (P0) or adult stage. *(B)* Heatmap of significant Gene Ontology (GO) terms (BP, biological processes; CC, cellular component; MF, molecular function) from GO-PCA analysis. Ten GO terms related to cytoskeleton changes or photoreceptor function and development are highlighted in bold. *(C)* UMAP visualization of regulon (i.e. a gene regulatory group consisting of a transcriptional factor and its downstream regulated genes) activities showing a separation of neonatal foveal and peripheral cones. *(D)* Waterfall plot showing the top ten regulons ranked by Regulon Specificity Score (RSS) in neonatal foveal and peripheral cones, respectively. *(E)* Fluorescence in situ hybridization validation of the expression of *SOX6, SOX4, and RAX* (green) in neonatal cones. Nuclei stained with DAPI are in magenta. Scale bar, 5 μm. *(F)* UMAP visualization of cell classes identified from scATAC-seq data. *(G)* Coverage plots showing peaks enriched at the loci of known cone marker genes, *GNAT2* and *OTX2*, across all cell classes. (H) UMAP visualization of foveal and peripheral cones from scATAC-seq data. (I) Dot plot showing the chromatin accessibility of selected TFs between foveal and peripheral cones. (J) Identification of significantly enriched binding motifs associated with differential accessible peaks between foveal and peripheral cones.

We used the Gene Ontology (GO)-PCA analysis (43) to find biological pathways that distinguished cones by age or region (Fig. 7*B*, *SI Appendix*, Fig. S9*E*). Neonatal foveal cones were enriched over neonatal peripheral cones, as well as all adult cones, in pathways associated with active morphogenesis such as “axon-dendritic transport”, “cytoskeleton-dependent intracellular transport”, and “protein localization to plasma membrane.” This result is consistent with the relatively immature state of foveal cones at birth. In adults, foveal and peripheral cones differed in pathways related to energy metabolism: adult foveal cones were enriched over all other groups in genes involved in glycolysis and gluconeogenesis, whereas adult peripheral cones were enriched over all other groups in pathways involved in oxidative phosphorylation, such as ATP coupled electron transport and proton-transporting ATP synthase complex. This difference raises the possibility that foveal and peripheral cones are powered in different ways (44).

Interestingly, the GO term “photoreceptor cell maintenance” identified multiple genes linked to retinitis pigmentosa (45) and Leber Congenital Amaurosis (LCA) (46) as selectively enriched in foveal cones (*SI Appendix*, Fig. S9*E*). Of note, retinal dehydrogenase 12 (*RDH12*), the causal gene for LCA13, was differentially expressed between foveal and peripheral cones at the neonatal stage, but the regional difference evened out by adulthood (SI Appendix, Fig. S9*E, F*). Although LCA13 affects the entire eye, macular dystrophy is an early and common feature (47). This result suggests that some genes associated with macular dystrophy may exert their effects at an early stage.

### Distinct transcriptional regulations in neonatal foveal and peripheral cones

Due to the distinct gene expression pattern of neonatal foveal cones, we used the single-cell regulatory network inference and clustering (SCENIC) algorithm to identify region-specific gene regulatory networks (GRNs) that could regulate their differentiation (48). This method generates selectively regulated gene sets called regulons that contain transcription factors (TFs) and their predicted target genes. We identified 219 regulons associated with 4,986 neonatal cones (*SI Appendix*, Fig. S10*A*). Regulon activity, like DEGs generally, separated neonatal cones into foveal and peripheral cohorts (Fig. 7*C*), demonstrating that some regulatory networks are differentially expressed in the two populations. The top regulons in each group are shown in Fig. 7*D*. The five regulons most enriched in foveal cones are AHR, PBX1, SOX6, BACH1 and FOXO3 as their defining transcription factors, while those most enriched in peripheral cones are defined by SOX4, RAX, LHX3, FOSB, and OLIG. Indeed, in many of these regulons, the key transcription factors and their target genes, whether positively or negatively regulated, show distinct enrichments between foveal and peripheral cones (*SI Appendix*, Fig. S10*B*). Hierarchical clustering of cones based on these five regulons confirmed a predominant separation by regions (*SI Appendix*, Fig. S10*C*). Notably, some of these TFs show differential expressions only at the neonatal stage, not in adults, suggesting their roles during the development (*SI Appendix*, Fig. S10*D*). Using fluorescence in situ hybridization, we confirmed the expression of SOX6 in foveal cones and the expressions of *SOX4* and *RAX* in peripheral cones at the neonatal retina (Fig. 7*E*, *SI Appendix*, Fig. S10*E*).

To verify the regulons inferred from scRNA-seq dataset, we generated single-cell ATAC-seq analyses of neonatal foveal and peripheral retina and identified open chromatin peaks associated with six main neuronal classes, including cones (Fig. 7*F, G*, *SI Appendix*, Fig. S10*F*). The clustering of all cones separated them into two clusters, representing foveal and peripheral cones, respectively (Fig. 7H). Comparing the chromatin accessibility associated with the TFs from the top regulons, we found that SOX6, AHR, and BACH1 show higher gene activities in foveal cones, while LHX3, FOSB, RAX and SOX4 have higher activities in peripheral cones, consistent with their regulon activities (Fig. 7*I*). We further determined the enriched motif binding sites of transcriptional factors (TFs) associated with differential peaks between foveal and peripheral cones. Among the TFs with significant binding motifs, six TFs are matched with top regulons (Fig. 7*J*). Altogether, these results verify most of the inferred regulons and demonstrate distinct transcriptional regulation in foveal and peripheral cones.

### Predicted interactions between foveal Müller glia and cones

We next turned to MG because, as noted above, they and cones show the greatest regional and developmental differences among cell types (Fig. 6, *SI Appendix*, Fig. S8*C* and *D*). This pattern suggests the possibility that MG maturation could promote cone maturation. Thus, we first investigated the potential biological functions associated with genes differentially expressed between neonatal foveal and peripheral MG (referred to as regional DEGs). By performing Protein-Protein Interaction (PPI) analysis of these regional DEGs, we can identify closely connected components via the Molecular Complex Detection (MCODE) algorithm, which aids in identifying functional modules within regional DEGs (see *SI Appendix*, Materials and Methods). Application of this method identified 13 components, annotation of which revealed 11 functional modules. Many modules are related to common cellular processes, such as oxidative phosphorylation, proteolysis, and mRNA processing. However, three modules – (1) Regulation of growth (*STAT1*, *INSR*, *FGF9*, *FGF13),* (2) Regulation of insulin-like growth factor (*IGFBP4*, *IGFBP5*, *SPP1),* and (3) collagen biosynthesis and extracellular matrix organization (*COL2A1*, *COL4A3*, *COL4A4*, *COL9A1*, *COL11A1)* – are of interest as they suggest a potential role of MG in morphogenesis (Fig. 8*A* and *C*). We validated expression of some of these regional DEGs by *in situ* hybridization (Fig. 8*B*).

**Figure 8.**
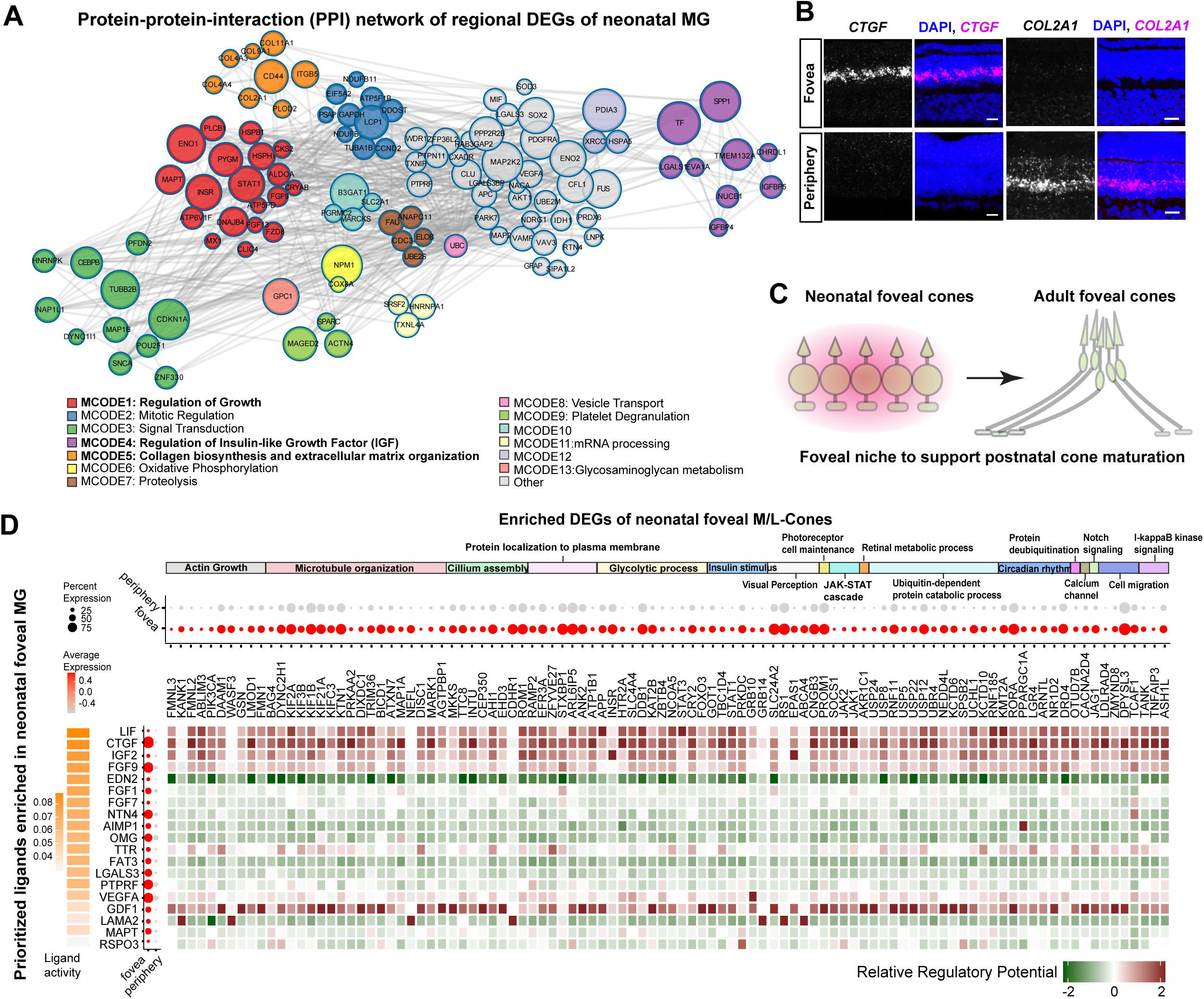
Foveal Müller glia ligands may regulate the expression of growth-related genes in adjacent foveal cones. *(A)* Protein-protein-interaction (PPI) network analysis of differentially expressed genes between neonatal foveal and peripheral Müller glia (MG) showing densely connected network components, which are involved in multiple growth-related GO terms. Each line indicates interactions between two proteins, while the circle size represents the connectivity degree, with larger circles indicating a higher degree. Distinct Molecular Complex Detection (MCODE) components are color coded, and annotations of MCODE components to functional modules are listed. “Other” means no MCODE complex detected. No functional modules are associated with MCODE 10 and 12. MCODE1, 4, and 5 are highlighted in bold. *(B)* Fluorescence in situ hybridization validations of *CTGF* (magenta) and *COL2A1* (magenta) expression in foveal and peripheral MG, respectively. Nuclei stained with DAPI are in blue. Scale bar, 20 μm. *(C)* A working model suggesting that the fovea provides a specific niche which enables the morphogenesis of postnatal cones. *(D)* NicheNet analysis predicting potential communication pathways between MG and cones. From left to right: orange heatmap showing the ranked ligand activity of nineteen ligands (secreted or membrane-bound) that are enriched in foveal MG; dot plot showing the expression of these ligands in the neonatal MGs; heatmap of the ligand-target interaction matrix denoting the relative regulatory potential between ligands from foveal MG and their target genes in foveal cones. From top to bottom: annotation bars grouping the target genes in foveal cones based on GO terms; dot plot showing the expression of these target genes in neonatal foveal cones.

In seeking MG-derived factors that could affect cones, we noted that several of the genes most enriched in neonatal foveal MG were secreted molecules known to interact with cellular receptors, including FGF9, FGF13, CTGF, VEGFA, SPP1 and several collagens. We therefore employed the computational method NicheNet (49) to predict the secreted ligands from MG that could influence the expression of genes enriched in foveal cones. First, we imputed all potential ligand-receptor pairs between foveal MG and foveal cones using the NicheNet prior model (*SI Appendix*, Fig. S11*A*). We then prioritized top active ligands based on their ligand activities, which are calculated by a statistical correlation between predicated expressions of target genes as downstream effects from each ligand-receptor interaction and the real expression levels of target genes. Among the 19 top-ranked ligands, we found five that broadly drive the expression of enriched genes in foveal cones, with many of these target genes being involved in cytoskeleton movement and cilium assembly (Fig. 8*D*). In contrast, when we used NicheNet to explore the opposite regulatory direction, namely secreted ligands from foveal cones that might influence expression of genes enriched in foveal MG, we found weaker ligand activity of foveal cones compared to MG, and few cases in which cone-derived ligands were predicted to drive expression of MG genes (*SI Appendix*, Fig. S11*B* and *C*). This asymmetry suggests the possibility that proteins secreted from neonatal foveal MG may play a role in the maturation of foveal cones, thereby providing a foundation for future mechanistic studies of foveal maturation (Fig. 8*C*).

## Discussion

It is important to study the retina of non-human primates because they share features with humans that are absent from other model systems, such as mice. The fovea is one prominent primate-specific structure. In this context, marmosets are of growing interest because their small bodies and relatively short gestation period suit them for biomedical research generally and visual science in particular (16). Despite previous histological and physiological characterization of marmoset retinal cell types (18, 21), the overall cellular composition of the marmoset retina remains undefined. In this study, we utilized scRNA-seq to characterize the cell types in the fovea and peripheral retina of both adult and neonatal marmosets, and compared them to those of macaque and human. We identified 65 foveal and 68 peripheral types in neonates and 68 foveal and 56 peripheral types in adults. These cell types are transcriptomically related across regions, ages, and species, but each comparison revealed intriguing differences in gene expression. This study provides a cell atlas of the marmoset retina that will facilitate primate retinal research and enable further analysis of foveal evolution and maturation.

### Foveal evolution

Marmosets are considered a more primitive primate species than macaque and humans, as marmosets have a smaller and smoother-surfaced brain (50). We suspected that marmosets might have a simpler fovea than humans or macaques, with cell types similar to those in a hypothetical primate ancestor. However, our cross-primate comparisons suggest a different model. First, the marmoset contains a large diversity of cell types, no fewer than those reported in macaques or humans (Fig. 2, 3). Moreover, the transcriptomic profile of foveal cell types displayed a high degree of correspondence across marmosets, macaques, and humans (Fig. 3). Lastly, fractions of individual foveal cell types are comparable between these three species. These results indicate that a pool of diverse cell types may have been established when the foveated primate ancestor emerged 35-40 million years ago. Notably, foveal size is almost uniform across primates, despite a large variation in total retinal area from marmosets to humans (9). These conserved features suggest that the primate fovea shares a common blueprint, and that its conserved cellular and molecular composition is critical for constructing a functional foveal circuit to support high-acuity vision. Furthermore, the conserved nature of the marmoset fovea suggests it may be particularly valuable for the study of macular degeneration and dystrophy (7, 22).

Despite the high degree of conservation of the primate fovea, several species-specific differences exist. A subtle but critical difference is that the marmoset fovea contains the lowest fraction of midget RGCs (MGCs) among the three primate species (Fig. 3). This finding is consistent with the fact that the marmoset has fewer laminated parvocellular layers that receive projections from MGCs compared to the macaque, but has a well-defined lamination of the koniocellular layer where other RGCs innervate (51, 52). In the midget pathway, the fraction of ON MGCs and the BCs that innervate them (called invaginating midget BCs or IMBs) showed increased dominance from marmoset to macaque to human. However, this pattern was not observed for the OFF MGCs or the bipolars that innervate them (flat midget BCs or FMBs). Thus, the modification of ON and OFF midget pathways may have occurred via distinct evolutionary trajectories. Supporting this hypothesis, a recent study using serial block-face scanning electron microscopy identified S-cone specific FMB in the human retina and macaque, but not in marmoset (53). Thus, the OFF midget pathway gained new circuit connections for color vision in macaque and human, but not in marmoset (54). The transcriptomic identification of OFFx BCs in the macaque and human fovea but not in the marmoset is also intriguing. Transcriptomic mapping has confirmed that OFFx BC is a homolog of a mouse BC type (BC1B) (26, 35, 37), suggesting that it may have been lost in marmoset.

### Foveal development

The transcriptomic mapping of foveal and peripheral cell types at neonates and adults revealed a high level of correspondence in the marmoset retina (Fig. 5). This finding aligns with our previous discoveries in the human and macaque retina, demonstrating that the fovea and peripheral retina share a similar pool of cell types (26, 27, 37). However, this raises the question of what factors contribute to the regional specialization observed in the fovea. By using regional scores to calculate the transcriptomic divergence in corresponding cell types between the fovea and peripheral retina, we identified several key themes. First, nearly all retinal cell types are fully specified at birth in both the fovea and peripheral retina. Second, the level of regional transcriptomic difference is not uniform across cell types, but rather more prominent in a few of select cell types. Third, regional differences increased during postnatal development (Fig. 6*A*-*D*, *SI Appendix*, Fig. S8*C*). Fourth, the level of developmental change is not uniform across cell types, with only selected cell types undergoing drastic developmental changes (Fig. 6*E*-*H*, *SI Appendix*, Fig. S8*D*).

Interestingly, the top three cell types showing the greatest regional and developmental differences are foveal Müller glia, cones, and MGCs (Fig. 6*E* and *F*). These three cell types undergo active modifications that facilitate foveal maturation. For example, in foveal cones, enriched biological pathways include cytoskeleton modification and photoreceptor development (Fig. 7*B*), which align with forthcoming morphological changes of foveal cones and are hallmark events that lead to foveal maturation (Fig. 1*B*).

Gene expression differences between foveal and peripheral cells are likely influenced by their local environment and controlled by distinct transcriptional regulation programs. Notably, foveal and peripheral cones showed divergent transcriptional regulation at the neonatal age. Among the top enriched transcriptional factors (TFs), a pair of Sry-related box transcription factors – SOX6 and SOX4 – and their regulated downstream genes, are expressed in foveal and peripheral cones, respectively (Fig. 7*D*, *SI Appendix*, Fig. S10*B*). The distinct expressions and gene activities of SOX6 and SOX4 between foveal and peripheral cones are confirmed by fluorescence in situ hybridization and scATAC-seq, respectively (Fig. 7*E* and *I*). Notably, active motif activities of SOX4 were detected in peripheral cones (Fig. 7*J*). SOX4 has been shown to mediate retinal development, while the role of SOX6 in the retina is unknown (55, 56). Interestingly, the expression of SOX6 and SOX4 is transient in cones as their expression is not detected in adults (Fig. 7*E*, *SI Appendix*, Fig. S10*D*, *E*), pointing to the importance of the early postnatal period for the transcriptional programs that lead to foveal maturation. Defects in acquiring the specific transcriptional machinery could cause altered foveal development, as observed in pediatric patients with retinopathy of prematurity (57). Furthermore, some of the identified TFs’ functions align with characteristic features of the fovea. For example, the BACH1 regulatory network is enriched in neonatal foveal cones (Fig. 7*D,* I, *SI Appendix*, Fig. S10*B*, *C*). BACH1 is known to repress Wnt/β-Catenin signaling and angiogenesis, which may be critical for maintaining the avascular area around the fovea (58). Future studies on the roles of these TFs in neonatal foveal cone maturation may open new avenues for the application of foveal cone differentiation in translational settings.

### The role of Müller glia in foveal development

Regional heterogeneity of Müller glia (MG) has been observed in the chick retina. Specifically, chicken MG show transcriptomic heterogeneity associated with their cardinal positions in the retina (59). Particularly in the high acuity area (HAA), a rod-free zone in the chicken retina, MG express a high level of FGF8, CYP26C1, and CYP26A1 (60). The enriched expression of these three genes is important for maintaining a low level of retinoic acid, which inhibits the fate of rods but promotes cone fate in the HAA. Thus, MG at distinct retinal locations could maintain a specific environmental niche to guide regional neurogenesis and neuronal development.

Previous studies of the macaque and human retina have identified a significant degree of difference in gene expression between foveal and peripheral MG (26, 27, 61). In this study of the marmoset retina, MG are also the most divergent cell type between the fovea and peripheral retina (Fig. 6*A*-*D*, *SI Appendix*, Fig. S8*C*). Thus, MG maintain a distinct molecular profile in the primate fovea. Many of these genes enriched in foveal MG, such as FGF9, are shared across primates, suggesting a conserved role in supporting foveal function (*SI Appendix*, Fig. S9*D*). Protein-protein interaction analysis of DE genes in foveal MG identified secreted molecules with known regulatory functions in cell growth or modification of the extracellular matrix (Fig. 8*A*). We thus hypothesized a potential role of these secreted molecules in foveal cone morphogenesis. Cell-cell interaction analysis via NicheNet predicted a strong regulation of gene expression in foveal cones by secreted ligands, such as FGF9 and CTGF (Fig. 8*D*). FGF9 is particularly interesting in that FGF signaling mediates the development of the outer plexiform layer in the zebrafish retina (62), and FGF receptors have enriched expression in primate foveal cones (63). Future confirmation of the source of FGF9 and an *in vivo* examination of the function of FGF9 are necessary to determine the role of foveal MG in mediating the cone morphogenesis. Additionally, CTGF has been shown to promote cell migration via modifying extracellular matrix (64). Thus, it is intriguing to speculate that CTGF may play a role in mediating cone migration, which is a prominent feature of fovea formation.

Genes enriched in peripheral MG could potentially be of interest as well. For example, COL2A1 is a gene enriched in peripheral MG (Fig. 8*A* and *B*). Mutations in COL2A1 cause Stickler syndrome in patients. These patients also have foveal hypoplasia, which means their retinas lack a fovea (65). As COL2A1 encodes the alpha-1 chain of type II collagen, the reorganization of the cellular matrix by COL2A1 in the peripheral retina could be critical for postnatal foveal development.

## Materials and Methods

### Tissue Procurement and Sequencing Library Preparation

Adult and neonatal eyes were enucleated from deeply anesthetized male and female marmosets (around two-years-old) at the time of death. The marmoset tissue collection was approved by and in accordance with the guidelines for the care and use of animals at Massachusetts Institute of Technology Institutional Animal Care and Use Committee. Marmoset eyes were collected from animals that had reached the end of unrelated studies. No ocular or visual abnormalities were noted. After enucleation, eyes were immediately placed in ice-cold hibernate medium (BrainBits) before dissection. The anterior chamber and the vitreous were removed by a rapid hemisection, and the posterior eyecup was immersed in room-temperature Ames’ medium (Sigma, equilibrated with 95% O2/ 5% CO2 for at least 20 minutes). For details of dissection, see *SI Appendix*, Materials and Methods.

### Single-Cell/Nuclei RNA-seq Library Preparation and Data Analysis

Single-cell RNA-seq libraries were generated and analyzed by methods described previously (26), modified as detailed in *SI Appendix*, Materials and Methods. For details of tissue processing and preparation of single-nuclei RNA-sequencing libraries, see *SI Appendix*, Materials and Methods. Sequencing reads (scRNA-seq and snRNA-seq) were aligned to calJac4, and initial quality control and quantification were performed using the Cell Ranger software (10X Genomics). For details of data processing and analysis, see *SI Appendix*, Materials and Methods.

### Single-Cell ATAC-seq Library Preparation and Data Analysis

Foveal and peripheral retinas from neonatal marmoset eyes were dissected and immediately snap-frozen using dry ice. The frozen tissues were stored at -80 °C freezer until the procedure. For details of library preparation, data processing, and analysis, see *SI Appendix*, Materials and Methods.

### Fluorescent In Situ Hybridization (FISH) validations and Image Processing

Marmoset eyes were fixed in 4% PFA, and then the anterior chamber was removed, following the procedure described under “Tissue Procurement.” The posterior poles containing the retina were rinsed with PBS and immersed in 30% sucrose at 4°C overnight, and then embedded with Tissue Freezing Medium (Electron Microscopy Sciences). The tissue was sectioned at 20 μm thickness and stored at -80°C for long-term storage. Fluorescent in situ probes against specific marmoset genes were generated following previously described methods (26), modified as detailed in *SI Appendix*, Materials and Methods. For details of image acquisition and processing, see *SI Appendix*, Materials and Methods.

## Data Availability

All the datasets were deposited to Gene Expression Omnibus (GEO: GSE249004).

## Code Availability

https://github.com/PengYRLab/MarmosetRetinalCellAtlas

## Acknowledgements

This work was supported by departmental startup funds from the UCLA, a career development award to Y.R.P. from Research to Prevent Blindness, a career starter award to Y.R.P. from Knights Templar Foundation, Klingenstein-Simons neuroscience fellowship (Y.R.P.), and an unrestricted grant to the Department of Ophthalmology from Research to Prevent Blindness, and NIH grants EY028633 and EY022073 (J.R.S.). The marmoset resource was supported by the Yang-Tan Collective at MIT, Tan-Yang Center for Autism Research at MIT, the Stanley Center for Psychiatric Research at Broad Institute of MIT and Harvard. We thank Raneesh Ramarapu for assisting with initial data analysis and technical assistant.

## Author Contributions

J.R.S and Y.R.P. conceived and supervised the study. L.Z., J.W., R.X., and Y.R.P. performed the computational analysis. Y.R.P. and M.C. performed sc/snRNA-seq, scATAC-seq, and histology experiments, Q. Z., and G. F. collected tissues. J.R.S and Y.R. P. wrote the paper with inputs from other authors.

## Corresponding authors

Correspondence to Yi-Rong Peng

