## Supplementary material for "Evolutionary and Developmental Specialization of Foveal Cell Types in the Marmoset": Supplmental figures

**Supplementary Information**

#### SI MATERIALS & METHODS

##### Tissue Procurement
The posterior eyecup was immersed in room-temperature Ames' medium and the retinal tissue was then separated from the sclera. The fovea (~1.5mm in diameter) and the peripheral retina (any area 5mm away from the fovea), were isolated for further preparation for scRNA-seq experiments. For the adult retinal tissue used for histology or fluorescence in situ hybridization (FISH), adult animals were perfused with 4% paraformaldehyde (PFA) under deep anesthesia. Eyes were then collected, post-fixed for 1 hour in 4% PFA, and stored in ice-cold Ames' solution. For the neonatal retina tissue used for histology or FISH, enucleated eyes were fixed for 1 hour in 4% PFA and then post-fixed for another 1 hour in 4% PFA after the anterior chamber was removed.

##### Single Cell Isolation and Single Cell RNA-Sequencing
Foveal and peripheral retinal tissues were dissected from neonatal and adult marmoset retinas. The foveal piece (~1.5mm in diameter), which contained the whole fovea, was dissected out and dissociated into individual cells for high throughput droplet-based single cell RNA sequencing (scRNA-seq, 10x Genomics). The peripheral area (any area 5mm away from the foveal center) was also dissociated into a single cell suspension and underwent two enrichment procedures. We first used CD73, a photoreceptor marker, to label photoreceptors and used a microbeads-conjugated secondary antibody to deplete all the CD73 labeled cells (1). Second, we enriched RGCs and ACs with CD90 positive selection via fluorescence-activated cell sorting (FACS). The peripheral cells after CD73 depletion or CD90 enrichment were used to generate scRNA-seq libraries (10X Genomics). Libraries were sequenced on the Illumina HiSeq 2500 platform.

##### Single Nuclei Isolation and Single Nuclei RNA-sequencing
The peripheral retinal tissues of an adult marmoset were snap frozen with dry ice. Nuclei were isolated with ice-cold Nuclei EZ lysis buffer and resuspended with 1%BSA/1X PBS. Nuclei were stained with anti-NeuN-PE (Millipore) for 30min at 4C. After washing and resuspension, the nuclei solution was additionally stained with Dyecycle Ruby (Thermo Fisher) before sorting. NeuN-positive and Dyecycle-positive nuclei were sorted via FACS. The sorted nuclei were used to generate single nuclei sequencing (snRNA-seq) libraries (10X Genomics) and were sequenced on the Illumina NovaSeq S4 platform.

##### Single-Cell RNA-Seq Data Analysis

###### *Read Alignment and Generation of Count Matrices*
Sequencing reads (scRNA-seq and snRNA-seq) were aligned to calJac4, and initial quality control and quantification were performed using the Cell Ranger software (10X Genomics). For mapping snRNA-seq reads, the "--include-introns--" mode was used to increase the mapping of reads from premature transcripts in nuclei. After read alignment, count matrices were generated for the snRNA-seq library and individual scRNA-seq libraries. For scRNA-seq data, count matrices were aggregated for neonatal and adult retinal cells, respectively, to create two separate datasets.

###### *Data Processing, Batch Correction, and Clustering*
We used the Seurat R package (Version 4) (2) for all the data processing and clustering steps. For each dataset, the count matrix was filtered based on a minimum of 400 expressed genes per cell and a minimum of 10 cells with detected expression per gene. We also filtered out cells with a high percentage of mitochondrial and ribosomal gene expression. The filtered data were

normalized using the “NormalizedData” function with a scale.factor of 10,000. Next, we detected highly variable features in the dataset using the “FindVariableFeatures” function with the “vst” method. Subsequently, we scaled the dataset based on the percentages of mitochondrial and ribosomal gene expressions, and the total number of transcripts detected per cell. After scaling, we performed the PCA analysis and used the Jackstraw plot to determine the number of significant principal components (PC) for clustering.

To correct biases associated with biological or technical replicates, we employed several batch correction methods, including Harmony (3), FastMNN, limma (4), and LIGER (5). We evaluate the performances of each method using the Cell-specific Mixing Score (CMS) (6) and the Local Density Differences (ldfDiff) score, and found that Harmony yielded the best results. Thus, we used Harmony for batch correction. Following batch correction, we used the “FindNeighbors” and “FindClusters” functions to determine cell clusters, which were then visualized in two dimensions using UMAP.

***Annotation of Cell Types in Individual Cell Classes***
Based on the expression of canonical marker genes for individual cell classes (1, 7), we assigned each cluster to one of the six cell classes and divided the dataset into six subsets corresponding to individual cell classes. For each cell-class subset, we further separated foveal and peripheral cells, resulting in twelve individual datasets for subsequent clustering. In the individual datasets, we filtered cells based on the quality and their specificity to their assigned cell classes. We then performed PCA analysis, batch correction with Harmony, clustering analysis, and cluster visualization using UMAP. We used the expression of maker genes previously found in the macaque dataset as a reference to assign cell types (1). For adult peripheral HCs and RGCs, we integrated snRNA-seq data with scRNA-seq data to identify the final cell types (see below).

**Single-Nuclei RNA-seq Data Processing, Clustering, and Annotation**
Similar to the analysis procedure for scRNA-seq data, the snRNA-seq dataset was firstly filtered based on a minimum of 200 genes per nucleus and a minimum of three nuclei per gene, resulting in a total of 3,127 nuclei. We further filtered out low-quality nuclei and potential doublets based on the numbers of transcripts and genes detected, as well as the percentages of mitochondrial and ribosomal gene expressions. Next, we performed data normalization, scaling, PCA analysis, clustering analysis, and then UMAP visualization. Each cluster identified was then assigned a cell class based on canonical marker genes. Due to the NeuN enrichment methods, we found ACs (~40% of total nuclei) and RGCs (~18%) were the two most abundant cell classes in the dataset. Additionally, we detected some HCs (~3%). Subsequently, we extracted RGCs and HCs for integration analysis with scRNA-seq data, aiming to improve the cell-type identification within the two cell classes.

**Integration of SnRNA-seq and ScRNA-seq Data for Clustering Adult Peripheral RGC and HC Cell Types**
We used canonical correlation analysis (CCA) (8, 9) to integrate the scRNA-seq and snRNA-seq data for adult peripheral RGCs and HCs, respectively, and performed the clustering analysis. Specifically, we first employed the “RunCCA” function with default parameters to construct a merged object containing both datasets. We then executed the standard Seurat clustering steps: NormalizeData, ScaleData, FindNeighbors (reduction=”cca”), FindClusters and RunUMAP (reduction=”cca”). After the analysis, the scRNA-seq and snRNA-seq data were effectively combined within individual clusters in the UMAP visualization (*SI Appendix*, Fig. S1*D*), and all the clusters identified in the integrated UMAP agreed well with clusters identified from each dataset.

**Clustering Foveal and Peripheral Cell Types Globally**

Firstly, we merged clusters from all the cell classes identified in adult fovea, adult periphery, neonatal fovea, neonatal periphery, respectively, resulting in four objects. In order to identify all the cell types in each object, we developed an iterative procedure to optimize the PC number and resolution value. Specifically, we initiated the process with a relatively low PC number and resolution value to obtain an initial clustering of all cells. Subsequently, we iteratively increased the PC number and resolution to enhance the cluster number. To evaluate the clustering stability of all the parameters tested, we constructed Clustering Trees (https://github.com/lazappi/clustree) (10) to determine the optimal PC and resolution value that would yield the maximum number of clusters while maintaining high stability. Consequently, we determined the use of a PC number of 300 and a resolution of 20 to maximize the clustering and generated a global UMAP of all clusters. To assign cell types for each cluster in the global UMAP, we conducted transcriptomic mapping to find the best-matched cell type from the individual cell-class UMAP to each cluster in the global UMAP.

The same analysis methods were used to integrate and cluster foveal and peripheral cell types, as well as neonatal and adult cell types, as shown in Figure 6*A*, *C*, *E*, *F*.

**Identification of Marker Genes for Individual Cell Types**

To identify cell-type–specific marker genes for individual cell types in each global UMAP, we first determined the differential expression genes (DEGs) for individual cell types using the "FindAllMarkers" function with the "MAST" test. In order to select the most specific DEGs for each cell type, we ranked the individual DEGs for each cell type based on the effective size, which was determined by the ratio of the expression percentage of one DEG in the queried cell type to that in all other cell types. After selecting the top three DEGs for all cell types, we plotted their expressions using the dotplot (Fig. 2*C* and 4*C*, *SI Appendix*, Fig. S2, 3, 6 and 7).

**Transcriptomic Mapping Across Regions, Developmental Stages, and Species**

We constructed a multi-class classifier using XGboost, as previously described (1), to establish correlations between cell types across regions, developmental stages, and species. Briefly, we first identified shared highly variable genes (HVGs) between the training and testing objects. We then extracted the normalized data matrix subsets containing the HVGs from these objects and used them as training and testing dataset, respectively. To train an XGboost model (11), we utilized the cell-type labels from the training object as classifier labels and used 60% of the cells (up to a maximum of 300 cells per type) from the training dataset to build the model. The model's performance was assessed by testing its prediction accuracy using the remaining training dataset. Once high-accuracy predictions were confirmed, we employed the model to predict the testing data matrix. Each testing cell was assigned to a training label, and the number of cells in each testing cluster assigned to individual training labels was summarized in a confusion matrix. Subsequently, we visualized the resulting confusion matrix by a Chord Diagram.

To map marmoset cell types to either macaque or human cell types, we followed the same procedure. For each cell type in individual marmoset classes, we constructed two separate XGBoost models using either macaque or human cell types as training data. We then classified the test marmoset cell types of the same cell class using these two models. For each marmoset cell, we define its mapped cell type to either macaque or human types as the one with higher prediction probability. We summarized the resulting confusion matrix into mapped proportions for each cell type and visualized it using a Chord Diagram, retaining the best-mapped cell type from either species for each marmoset cell type.

**Transcriptomic Comparison of Foveal Cell Types Among Three Primate Species**

##### *Integrating Foveal Cells from Three Primate Species, Clustering, and Generating A Dendrogram*

We pooled 200 cells per type from marmoset, macaque (1), and human (7) and used CCA to integrate the data. We identified 26 cell clusters with 100 PCs and resolution 3. Based on the expression of marker genes, we assigned these 26 clusters to 26 cell types/subclasses, each including cells from all three species. We then used the "BuildClusterTree" function to generate a dendrogram of cell types among the three species.

##### *Identification of Species-Conserved and -Specific Markers*

To identify conserved markers for homologous cell types among primates, we conducted differential expression gene (DEG) analysis for each homologous cell type using the "FindConservedMarkers" function from the Seurat R package. For cell types where specific genes could not be identified using this method, we used the "FindMarkers" function to identify a new set of markers and checked their cross-species conservation. We also examined the conservation of cell-type specific markers from the macaque atlas (1) to further refine the marker choices.

To identify cell-type–specific marker genes that are unique to each primate species, we used the "FindAllMarkers" function from the Seurat Package on each cell type within each species, comparing it against all other cells with a log fold change threshold of 1. For each identified gene, we calculated its effective size as the ratio between the percentage of the gene expressed in the current cell type and that in the remaining types. We selected the top 20 genes with the highest effective size for each type and further narrowed the selection down to those with an average log2 fold change greater than 1.5 and a percentage expression in the remaining cells of less than 0.1.

##### *Calculate the Proportions of Cell Types in Individual Cell Classes*

To compare the composition of foveal cell types across the three primate species, we calculated the proportion of each homologous cell type within its cell class using the original datasets (Marmoset: this study; Macaque: Peng et al., 2019; Human: Yan et al., 2020). We included an additional dataset (12) to calculate the proportions of human photoreceptors as well (*SI Appendix*, Table S3).

Finally, we assembled the following information: (1) the integrated UMAP, (2) the cell-type dendrogram, (3) the dotplot of conserved markers, and (4) the barplot of cell type proportions from inner to outer layers, into a Circos plot using the circlize R package (Fig. 3*A*).

##### Transcriptomic Correlations of Foveal Cell Types Among Three Primate Species

To determine transcriptomic convergence of homologous foveal cell types among the three primate species, we first merged 29,160 adult marmoset foveal cells, 92,626 adult macaque foveal cells (1), and 55,108 adult human foveal cells (7). Secondly, we used reciprocal PCA (RPCA) implemented in the Seurat R Package for data integration. We then performed unsupervised clustering and identified homologous cell types. Finally, we obtained the pseudobulk transcriptional profiles for each cell type in each species using the top 2,000 highly variable genes (HVGs). For each homologous type, we calculated the *Pearson* correlation coefficient of the average expression of HVGs between each pair of the three species (Fig. 3*B*).

##### Signed Gene Set Enrichment Analysis (sGSEA)

We developed a signed z-score-based gene set enrichment analysis (GSEA) to address the cellular changes during development. This method differs from conventional GSEA (13-15) as it considers the distinct responses (upregulation or downregulation) of genes within the gene set. The sign (positive/negative) of the change is considered in this method.

The normalized enrichment score (NES) is given by the integrated z-scores ($Z_{Stouffer}$) of the genes in the differentially expressed gene set using Stouffer's method.

For a given set of DEGs, the mean and standard deviation of DEG expression in the control group are first calculated. Then, the z-score of the *i*th DEG in cell *j* of the query group is given by:

$$z_{ij} = \frac{x_{ij} - mean(x_{i,reference})}{sd(x_{i,reference})}$$

where $x$ is the normalized gene expression value.

Next, the Stouffer method is employed to integrate the z-scores. The z-scores are weighted by the log2FC. The weighted Stouffer z-score is given by:

$$Z_{Stouffer} = \frac{\sum_{i \in Set} w_i z_i}{\sqrt{\sum_{i \in Set} w_i^2}}$$

where gene $i$ belongs to the gene set comprising both top up-regulated and down-regulated genes, ranked by p-value, weight $w_i$ is the signed average log2FC of the gene $i$, and $z_i$ is the z-score of gene $i$.

The weight $w_i$ is given by:

$$w_i = \begin{cases} log2FC, & i \in up - regulated \\ -log2FC, & i \in down - regulated \end{cases}$$

Under the null hypothesis, $Z_{Stouffer}$ follows the standard normal distribution:

$$Z_{stouffer} \sim N(0, 1)$$

Thus, the right-tailed p-value of significance can be derived directly by:

$$P = 1 - \Phi(Z_{stouffer})$$

where $\Phi$ is the cumulative distribution function of the standard normal distribution.

The R implementation of the algorithm is available at https://github.com/JunqiangWang/sGSEA.

**Earth Mover's Distance Calculation and Transformation**

The Earth mover's distance is a distance measure between two distributions (16, 17). Assume there are signature P and signature Q associated with partitions $P = \{(p_1, w_{1p}),\ (p_2, w_{2p}), \ldots, (p_n, w_{np})\}$ and $Q = \{(q_1,\ w_{1q}), (q_2,\ w_{2q}), \ldots,\ (q_m,\ w_{mq})\}$, where *p* and *q* are the partitions, and *w* is the amount of mass in the partition. We can find a flow F that solves the optimal transport problem:

$$min_F \langle c, f \rangle = min_F C_{i,j} F_{i,j}$$

subject to the constraints:

$$F \in R_+^{n \times m},\ \sum_j F_{i,j} \le w_{ip},\ \sum_i F_{ij} \le w_{jq} \text{ and } \sum_i \sum_j F_{i,j} = \min\{\sum_i w_{ip}, \sum_j w_{jq}\}$$

where $F_{ij}$ describes the amount of the mass flowing from $P_i$ to $Q_j$ and $C_{ij}$ describes the costs. The EMD is given by the normalized cost:

$$EMD(P, Q) = \frac{\sum_i \sum_j F_{i,j} C_{i,j}}{\sum_i \sum_j F_{i,j}}$$

We derived the EMD score to rank the regional differences or developmental changes among cell types. Firstly, we identified the differentially expressed genes (DEGs) for each cell type using the "FindMarkers function" with the MAST test (log2FC>1, p_val_adj<0.05) (18). We then used the "calculate_emd_gene" function of the EMDomics R package (19) to calculate the EMD distance using the aggregated expression of all DEGs for each corresponding type between the foveal and peripheral regions or between the adult and neonatal stages. To better visualize the changed level in each cell type relative to the mean change in all cell types, EMD values were then transformed into z scores in the final barplots (Fig. 6*B*, *D*, *F* and *G*).

**GO-PCA Analysis of M/L-Cones**

We used GO-PCA (https://github.com/flo-compbio/gopca) (20) to identify significantly enriched Gene Ontology (GO) terms with top principal components among the cone dataset (Fig. 5*A*). Briefly, cones within each type were randomly aggregated into multiple instances with 20 cells per instance, and the resulting matrix was used as input for GO-PCA, which was run with default parameters. We generated signatures of 94 GO terms. After excluding quality-driven GO terms, such as apoptotic signal pathways and ribosomal gene pathways, we curated signatures of 28 GO terms. Furthermore, we down-sampled each cone type to 50 instances and generated the heatmap using the 28 GO terms (Fig. 5*B*).

**Single-Cell Regulatory Network Inference and Clustering (SCENIC) Analysis of Neonatal M/L-Cones**

We used pySCENIC (21) to infer the gene regulatory networks (GRNs) among neonatal cones. Firstly, we filtered the expression matrix of all neonatal cones by removing genes with less than 3 counts in 1% of cells. The filtered matrix was used as the input matrix for SCENIC. We curated all the transcription factors (TFs) in the matrix using the human TF list (https://github.com/aertslab/pySCENIC/blob/master/resources/hs_hgnc_tfs.txt) and inferred potential regulons by identifying TF-target co-expression networks using the GRNBoost2 algorithm (22). Indirect targets (without TF binding motif) for each regulon were trimmed out using RcisTarget (23), resulting in the final regulons. Finally, the activity of each regulon was calculated with AUCell for each cell. We then clustered the cells based on their regulon activities and visualized them in the UMAP. The UMAP showed a clear separation between foveal and peripheral cones, indicating distinct GRNs for cones from distinct regions. To identify the regulons that are specific to foveal or peripheral cones, we calculated the Regulon Specificity Scores (RSS) (24) for each regulon in foveal and peripheral cones, respectively, using the "regulon_specificity_scores" function. We ranked the regulon activity and highlighted the top ten regulons for each cone type (Fig. 7*D*).

**Protein-Protein Interaction Network Analysis for Neonatal Müller Glia**

We used the web-based user interface platform of Metascape (25) to identify associations among all the differentially expressed genes (DEGs) between neonatal foveal and peripheral MG and visualized the protein-protein interaction (PPI) network using Cytoscape (26). We then used a molecular complex detection (MCODE) algorithm to identity the densely connected network of protein-protein interactions. The resulting MCODE components were functionally annotated using GO terms, Reactome (https://reactome.org/content/detail/R-HSA-1643685), and Kyoto Encyclopedia of Genes and Genomes (KEGG) pathways. We then imported the MCODE_PPI.cys file into Cytoscape to enhance the visualization of individual protein names. To focus on strong connectivity, we pruned the original network with a minimum connectivity degree of four.

**NicheNet Analysis to Infer the Interaction between Neonatal MG and M/L-Cones**

We used NicheNet (27) to investigate whether MG-ligands might regulate the expression of relevant genes in adjacent foveal M/L-Cones via the following steps: (1) Determine the gene sets from the sender cells (MG) and receiver cells (cones), respectively. We used the "FindMarkers" Function with the "MAST" test to identify the upregulated genes in foveal MG compared to peripheral MG as the sender gene list. We also identified the upregulated genes in foveal cones compared to peripheral cones as the initial receiver gene list. To facilitate the identification of functional interactions that promote cones' growth and other biological processes, we annotated the functions of the initial receiver genes using DAVID (28). We selected genes associated with 22 GO terms that were relevant to biological functions of cones, such as photoreceptor cell outer segment organization (GO:0035845), non-motile cilium assembly (GO:1905515), and actin cytoskeleton organization (GO:0030036). This resulted in 109 genes as a final gene list for receiver cells. (2) Establish ligand-target interactions and regulatory strengths. Based on the two gene lists from sender and receiver cells, we identified all the ligand-target pairs with a data-driven prioritization nomination from the NicheNet. We ranked ligands based on their ligand activity and obtained the regulatory potential scores for interactions between the top ligands and all 109 target genes from the receiver cell. We further scaled the regulatory potential scores of ligands for each target gene to identify which ligand has the strongest regulatory potential (Fig. 8*D*). (3) Infer the ligand-receptor links. Firstly, we identified all potential receptors for the top ligands from the NicheNet prior model. Then, we selected receptors that showed expression in receiver cells. We extracted the interaction potential and generated the ligand-receptors links (*SI Appendix*, Fig. S11*A*).

We performed the same analysis using NicheNet to infer the interaction between foveal cones as sender cells and MG as receiving cells.

#### Single-cell ATAC-seq Library Preparation and Data Analysis

##### *Generate ScATAC-seq libraries*

Frozen foveal and peripheral retinas were retrieved from a -80 °C freezer and placed on ice. Nuclei from these frozen samples were separately isolated using Lysis Buffer and Nuclei Isolation Columns from the Chromium Nuclei Isolation Kit (10X Genomics) following the Manufacturer's User Guide. The isolated nuclei were then resuspended in Resuspension Buffer (10X Genomics). ScATAC-seq libraries were generated from the recovered nuclei using the Chromium Next GEM Single Cell ATAC Reagent v2 Kit (10X Genomics). The quantity and quality of the scATAC-seq libraries were evaluated using Qubit fluorometers (Thermo Fisher) and Tapestation (Agilent). ScATAC-seq libraries were sequenced on the Illumina Novaseq X Plus 10B platform.

##### *scATAC-Seq data analysis*

Read alignment. We used the cellranger-atac software (v2.1.0, 10X Genomics) to map scATAC-seq reads. We compared two marmoset genomes for the mapping: "mCalJa1.2.pat.X" (GCF_011100555.1) from NCBI and "mCalJac1.pat.X" (GCA_011100555.1) from Ensembl. We found the latter resulted in slightly better mapping metrics, such as higher fractions of high-quality fragments overlapping peaks, high-quality fragments in cells, transposition events in peaks in cells. We thus used the "mCalJac1.pat.X" genome to build the reference package and also used Ensembl based annotation packages "ensembldb" to facilitatee peak annotations for downstream workflow. More specifically, we used the "cellranger-atac mkref" command with a custom configuration file and JASPAR vertebrate non-redundant motifs file (https://jaspar.elixir.no/download/data/2022/CORE/JASPAR2022_CORE_vertebrates_non-redundant_pfms_jaspar.txt) to constructure the reference. We then mapped individual samples to this reference using the "cellranger-atac count" command and processed each sample separately for sample quality control and primary classification.

Pre-processing. We used the Signac (v1.9.0) (29) workflow and other packages for downstream analysis. Firstly, we built a Signac object using the output files from cellranger. For sample quality control (QC), in addition to quality metrics provided by cellranger, we further evaluated the proportion of peaks mapped to different genomic regions such as exon, intron, intergenic and promoter regions. Specifically, we extracted the genomic ranges of all peaks from the Signac object using the granges() function, which were then transformed and saved as a bed file. Secondly, we used the annotatePeaks.pl program (HOMER 4.11.1) in conjunction with the fastq and gtf file of "mCalJac1.pat.X," to associate peaks with nearby genes. To incorporate gene annotation information to the object, we used the GetGRangesFromEnsDb() function to extract gene annotations from EnsDb. As an ensembldb annotation package for mCalJac1.pat.X is not available on Bioconductor, we sourced the AH113598, Ensembl 110 EnDb (corresponding to mCalJac1.pat.X) for marmoset from the AnnotationHub package. Lastly, we computed QC metrics including transcriptional start site (TSS) enrichment score, nucleosome signal score per cell, fraction of reads in peaks, and examined the nucleosome banding pattern for each sample. Cells that were outliers in these QC metrics were removed.

Clustering and cell class identification in each sample. We applied a latent semantic indexing (LSI) workflow as follows. Firstly, we conducted term frequency-inverse document frequency (TF-IDF) normalization using the RunTFIDF() function in Signac. For dimensional reduction, we used all features from the scATAC-seq data and performed singular value decomposition (SVD) on the TD-IDF matrix using the RunSVD() function. We excluded the first LSI component as its strong association with sequencing depth. To cluster the cells, we used the RunUMAP(), FindNeighbors() and FindClusters() functions (reduction="lsi", dims=2:30) from the Seurat package.

To classify cell clusters, we quantified gene activity based on fragments intersecting the gene body and promoter region using the GeneActivity() function. We then performed log normalization on this gene activity matrix with a scale factor of the median nCount_RNA per cell. We classified the cells using two approaches: (1) examining the activities of canonical retinal cell class marker genes and (2) integrating with scRNA-seq data from neonatal marmosets based on shared correlation patterns in the gene activity matrix and the scRNA-seq data. For this purpose, we used classified scRNA-seq data from the neonatal marmoset retina, which includes all retina cell classes from both the fovea and periphery regions. We first used the FindTransferAnchors() function (reduction="cca") to identify anchors between the scATAC-seq and scRNA-seq data, which are pairs of mutually nearest neighbor cells. We then input these anchors into the TransferData() function (weight.reduction = object[['lsi']], dims = 2:30) to predict a cell type score for each cell from the scATAC-seq data. We found a good correspondence between the cell type classification based on marker gene expression from the imputed gene activity matrix and the prediction by label transfer from the scRNA-seq data, demonstrating the robustness of the imputed gene activity data. These approaches enabled the identification of all major retinal cell classes in foveal and peripheral scATAC-seq datasets.

Merging and classifying a combined scATAC-seq data. Since peak calling was performed separately on each dataset, the peaks differ between samples. To compare foveal and peripheral scATAC-seq data, we first created a common set of peaks. This involved extracting genomic ranges of each sample using the makeGRangesFromDataFrame() function, based on the sample peak bed file, and then merging intersecting peaks between samples using the reduce() function from the GenomicRanges package. Low-quality peaks were filtered out based on peak width, and the peaks in each dataset were quantified. Furthermore, low count cells were removed, and fragments objects were constructed using the CreateFragmentObject() function, followed by creating a peaks X cell matrix for each dataset using the FeatureMatrix() function. Using these matrices, we created a Seurat object for each sample with the fragment object embedded. Based on their common set of peaks, we then merge these objects into a combined object.

To classify this combined object, we followed a similar procedure as described above for normalization and dimension reduction, adding a step for batch correction across samples. We first applied RunTFIDF() for normalization post-merging to ensure consistent IDF weighting across datasets. Next, we performed all the features to apply singular value decomposition (SVD) on the TF-IDF matrix using the RunSVD() function. Afther that, batch correction on the "ATAC" assay was performed using RunHarmony() (group.by.vars="dataset",reduction.use="lsi") function. Iterative application of RunUMAP(), FindNeighbors(), and FindClusters() (reduction = "harmony"), along with filtering out outler cells/clusters based on QC metrics, such as nCount_ATAC and nucleosome_signal, continued until no low-quality cells remained. We classified the final clustering using the two approaches mentioned earlier. Additionally, we validated our classification by examining peak profiles near known retinal marker genes using the CoveragePlot() function. We observed enriched peak signals around the promoter/enhancer regions of these genes, supporting our classification.

Clustering scATAC-seq data for foveal and peripheral cones. To compare the chromatin accessibility profiles between foveal and peripheral cones, we subsetted all cones from the combined object of all cell classes. We clustered all cones using a similar procedure without batch correction by harmony. The resulting UMAP showed distinct clusters for foveal and peripheral cones, indicating their unique epigenetic landscapes.

Motif analysis. We used the scATAC-seq data of all cones to derive top motifs and their associated transcription factors in the foveal and peripheral cones. To this end, we first curated a list of motif position frequency matrices (pfm) from the JASPAR database using the getMatrixSet() function (collection = "CORE", tax_group = 'vertebrates'). We then prepared a BSgenome genome package of mCalJac1.pat.X, required by the Signac AddMotifs() function. This involved downloading compressed fasta files of each chromosome of mCalJac1.pat.X from Ensembl, preparing the BSgenome data package seed file, and using the forgeBSgenomeDataPkg() function from the BSgenome package to create the source tree of the target package. We used the built BSgenome.mCalJac1.pat.X and the pfm obtained above as input for the AddMotifs() function to add motif information to our object. We then identified differentially accessible peaks between the foveal and peripheral cones, using the FindMarkers() function (test.use = 'LR',min.pct = 0.05, latent.vars = 'nCount_peaks'). This enabled us to identify top differentially accessible peaks. We performed hypergeometric test to identify enriched motifs in these differential peaks using the FindMotifs() function. Finally, we visualized the position weight matrices for the motifs using the MotifPlot() function.

**Fluorescent In Situ Hybridization (FISH) validations**

Fluorescent in situ probes against the marmoset genes *CTGF, COL2A1*, *RDH12*, and *OPN1MLW* were generated following previously described methods (1). Briefly, total RNA was extracted from marmoset retinas and converted to cDNA libraries through reverse transcription using the AzuraQuant cDNA synthesis kit. Antisense probe templates for individual target genes were PCR-amplified from the cDNA libraries using a reverse primer with a T7 sequence adaptor to permit in vitro transcription. DIG rUTP (Roche) and Fluorescein rUTP (Roche) were used to synthesize probes for single or double FISH experiments. Retinal sections were thawed, treated with 1.5 μg/mL of proteinase K (NEB), post-fixed with PFA, and deacetylated with acetic anhydride. After blocking, the retinal sections were incubated with probes overnight. Probe detection was performed with anti-DIG HRP (1:1000) and anti-Fluorescein POD (1:1000) followed by tyramide amplification (30).

HCR split-initiator probe sets against the marmoset genes *SOX4*, *SOX6*, and *RAX* were used for HCR RNA-FISH (31). Probe sets were synthesized by Molecular Instruments (www.molecularinstruments.com) using the coding sequence regions of the following NCBI Reference Sequences: XM_035294784.2 (SOX4), XM_035264180.2 (SOX6), and XM_002757287.5 (RAX). Retinal sections were fixed in 4% PFA for 15 minutes at 4°C and then sequentially immersed in 50% EtOH, 70% EtOH, 100% EtOH, and fresh 100% EtOH for five minutes each at room temperature. Slides were then immersed in three sequential washes with 1x PBS before proceeding to the HCR assay. Sections were incubated with probe hybridization buffer (Molecular Instruments) at 37°C for 10 minutes prior to overnight incubation with probe solution consisting of 0.4 pmol probe set in probe hybridization buffer (Molecular Instruments) at 37°C. Excess probes were removed by incubating the slides for 15 minutes at 37°C in each: 25%, 50%, 75%, and 100% 5x saline sodium citrate with 0.1% Tween 20 (SSCT) in probe wash buffer (Molecular Instruments). Slides were then immersed in 5x SSCT for five minutes at room temperature before proceeding to the amplification stage. Sections were pre-amplified with amplification buffer (Molecular Instruments) for 30 minutes at room temperature and then incubated overnight in a hairpin solution consisting of 6 pmol each of previously snap-cooled hairpins h1 and h2 (Molecular Instruments) in amplification buffer. Excess hairpins were removed by incubating slides in room temperature 5x SSCT twice for 30 minutes and once for five minutes. Sections were stained with DAPI (1:1000) in 5x SSCT for 20 minutes and cover slipped in antifade mounting reagent.

**Image Acquisition, Processing, and Analysis**

Images were acquired on an Olympus FluoViewTM FV1000 confocal microscope with 405, 488, and 599 lasers and scanned with 40X or 60X oil objective at the resolution of 1024x1024 pixels, a step size of 1 mm, and an 80 mm pinhole size. Maximum intensity projections were generated using ImageJ (NIH) software, and brightness and contrast adjustment were made using Adobe Photoshop CC.

#### Supplementary Figure Legends

**Fig. S1.** Data quality of the cell atlas of the adult marmoset retina. (*A*) Violin plots showing distributions of the number of expressed genes (nFeature_RNA), RNA counts, and percentages of mitochondrial genes (percent.mt) and ribosomal genes (percent.ribo) detected in the foveal and peripheral samples. (*B*) Uniform Manifold Approximation and Projection (UMAP) of all adult foveal cells, colored by individual sample identities. (*C*) UMAP visualization of 65 cell types in the fovea. (*D*) UMAP visualization of peripheral RGCs, as shown in Fig. 2B (RGC), but colored by different methods – scRNA-seq or snRNA-seq. (*E*) UMAP visualization of 53 cell types in the peripheral retina.

**Fig. S2.** Dot plot showing the expressions of marker genes for individual foveal cell classes (bottom 15 rows) and types (remaining rows) in the adult marmoset retina.

**Fig. S3.** Dot plot showing the expressions of marker genes for individual peripheral cell classes (bottom 15 rows) and types (remaining rows) in the adult marmoset retina.

**Fig. S4.** Comparison of foveal cell types from human, macaque, and marmoset. (*A*) UMAP visualization of 8 subclasses from 23,966 cells integrated from three species, same as shown in Fig. 3A. (*B*) UMAP visualization of 26 cell types/subclasses from 23,966 cells. (*C*) UMAP visualization of 23,966 cells colored by species, showing all clusters consist of cells from all three species. (*D*) Dot plot showing species-specific marker genes that are expressed by individual homologous cell types. (*E*) Chord diagrams presenting confusion matrices to show the transcriptomic mapping of marmoset foveal cells to their better matched cells between human and macaque. From left to right, the highest diversified cell classes–BCs, RGCs, and ACs, were used for the comparison. Each line from the marmoset cells points to its better matched counterpart between human and macaque cells.

**Fig. S5.** Data quality of the cell atlas of the neonatal marmoset retina. (*A*) Violin plots showing distributions of the number of expressed genes (nFeature_RNA), RNA counts, and percentages of mitochondrial genes (percent.mt) and ribosomal genes (percent.ribo) detected in the foveal and peripheral samples. (*B*) UMAP of all neonatal foveal cells, colored by individual sample identities. (*C*) UMAP of 62 cell types in the fovea. (*D*) UMAP of all neonatal peripheral cells, colored by enrichment methods: CD90+ (enrich RGCs and ACs) or CD73+ (deplete rods). Most of the RGCs were detected from the CD90+ sample, validating the efficacy of the enrichment methods. (*E*) UMAP of 66 cell types in the peripheral retina.

**Fig. S6.** Dot plot showing the expressions of marker genes for individual foveal cell classes (bottom 15 rows) and types (remaining rows) in the neonatal marmoset retina.

**Fig. S7.** Dot plot showing the expressions of marker genes for individual peripheral cell classes (bottom 15 rows) and types (remaining rows) in the neonatal marmoset retina.

**Fig. S8.** Comparison of foveal cell types across regions and across stages. (*A*) Pearson correlations of normalized enrichment score (NES) in individual cells between using 200 and using100 differentially expressed genes (DEGs). (*B*) Pearson correlations of NES in individual cells between using 200 and using 300 DEGs. The high correlation in both *A* and *B* demonstrated that the sGSEA method provides consistent results regardless of the number of DEGs used. (*C*) Box and whisker plot showing the quartiles of -log10FDR of regional scores from all the integrated neonatal (with suffix "_N") and adult cell types (with suffix "_A"), which were ranked from highest to lowest. FDR: false discovery rate. (*D*) Box and whisker plot showing the quartiles of -log10FDR of developmental scores from all the foveal (with suffix "_F") and peripheral (with suffix "_P") cell types, which were ranked from highest to lowest. (*E*) Examples of calculations of Earth mover's distance (EMD) in Figure 6B, D, F, and H. Density distribution of DEG expression levels for adult (pink) and neonatal (grey) cell populations of foveal MG. (*F*) Density distribution of DEG expression levels for adult (pink) and neonatal (grey) cell populations of foveal IMB.

**Fig. S9.** Comparison of foveal enriched genes across three primates and fovea-specific GO-PCA signatures in the marmoset. (*A*, *C*) Venn diagrams showing the numbers of genes that exhibited at least a two-fold enrichment (log2FC > 1) in foveal cones (*A*) and MG (*C*) compared to their peripheral counterparts, for individual primate species. This diagram also illustrates the numbers shared between each pair of species and those common to all three. (*B*, *D*) Waterfall plots showing DEGs between foveal and peripheral M/L-cones (*B*) and MG (*D*) in the marmoset, highlighting DEGs shared with the other two primates (left) and DEGs specific to the marmoset (right). The names of shared DEGs with an average log2 fold change (log2FC) greater than 2 or less than -2 are highlighted in green. The names of marmoset-specific DEGs with an average log2FC greater than 2 and less than -4 are lighted in orange. (*E*) Heatmap of expression patterns for selected GO signatures. (*F*) Double fluorescence in situ hybridization of *RDH12* (magenta) and *OPN1MLW* (green) to validate that the difference in expression levels of *RDH12* between foveal and peripheral cones is only observed at the neonatal stage, and not in the adult retina. Nuclei stained with DAPI are shown in blue. Scale bar, 20 μm. In each RDH12 image, the area containing photoreceptor segments is demarcated with a red bar and is magnified below. Scale bar, 20 μm.

**Fig. S10.** SCENIC analysis and scATAC-seq data. (A) Hierarchical clustering heatmap showing the binary regulon activity matrix of all the regulons identified from all neonatal cones. Cells tend to be aggregated by their region identity based on regulon activities. Blue boxes highlight top regulons in foveal cones. Orange boxes highlight top regulons in peripheral cones. *(B)* Heatmap of expression patterns of genes in regulons enriched in neonatal foveal and peripheral cones, respectively. *(C)* Heatmap of expression for the top five regulons among neonatal cones. Hierarchical clustering of cells (rows) and regulons (columns) showing two modules with a large separation of foveal and peripheral cones and distinct regulons associated with each type. (*D*) Dot plots showing the expression patterns of BACH1, SOX6, RAX, and SOX4 in cones of neonatal marmosets, adult marmosets, and adult macaques. *(E)* Fluorescence in situ hybridization validation of the expression of *SOX6, SOX4, and RAX* (green) in the central and peripheral retina. Nuclei are stained with DAPI (magenta). Scale bar, 20 μm. The area containing the cone layer is demarcated with a dashed box and is magnified below. Scale bar, 5 μm. The magnified cone layer is also shown in Fig. 7E. (*F*) UMAP visualization of cells from scATAC-seq data, as shown in Fig. 7F, but colored by different samples.

**Fig. S11.** Potential ligand-receptor interactions between MG and cones. (*A*) Heatmap showing potential receptors in foveal cones, which could bind to the top 19 ligands from foveal MG. Dot plots show the expression levels of detected ligands and receptors in foveal MG and cones, respectively. (*B*) Histogram comparison of top ligand activities between foveal cones (blue) and foveal MG (red). The ligand activity in cones is much lower than that in MG, indicating a lower likelihood of a regulatory relationship from cones to MG. (*C*) Heatmap showing the regulatory potential from ligands in foveal cones to target genes in foveal MG, supporting a weak regulatory relationship in this direction.

**Supplementary Table Information**

**Table S1.** Cell number and types in each dataset.

**Table S2.** Abbreviations for cell classes and types.

**Table S3.** Proportions of foveal cell types across three primates.

**Table S4.** The correspondence between cell types in Figure 5.

**Table S5.** The correspondence between cell types in integrated UMAPs and those in individual UMAPs, as related to Figure 6.

**Table S6.** The correspondence between cell types in integrated UMAPs and those in individual UMAPs, as related to Figure 6.

#### Figure S1

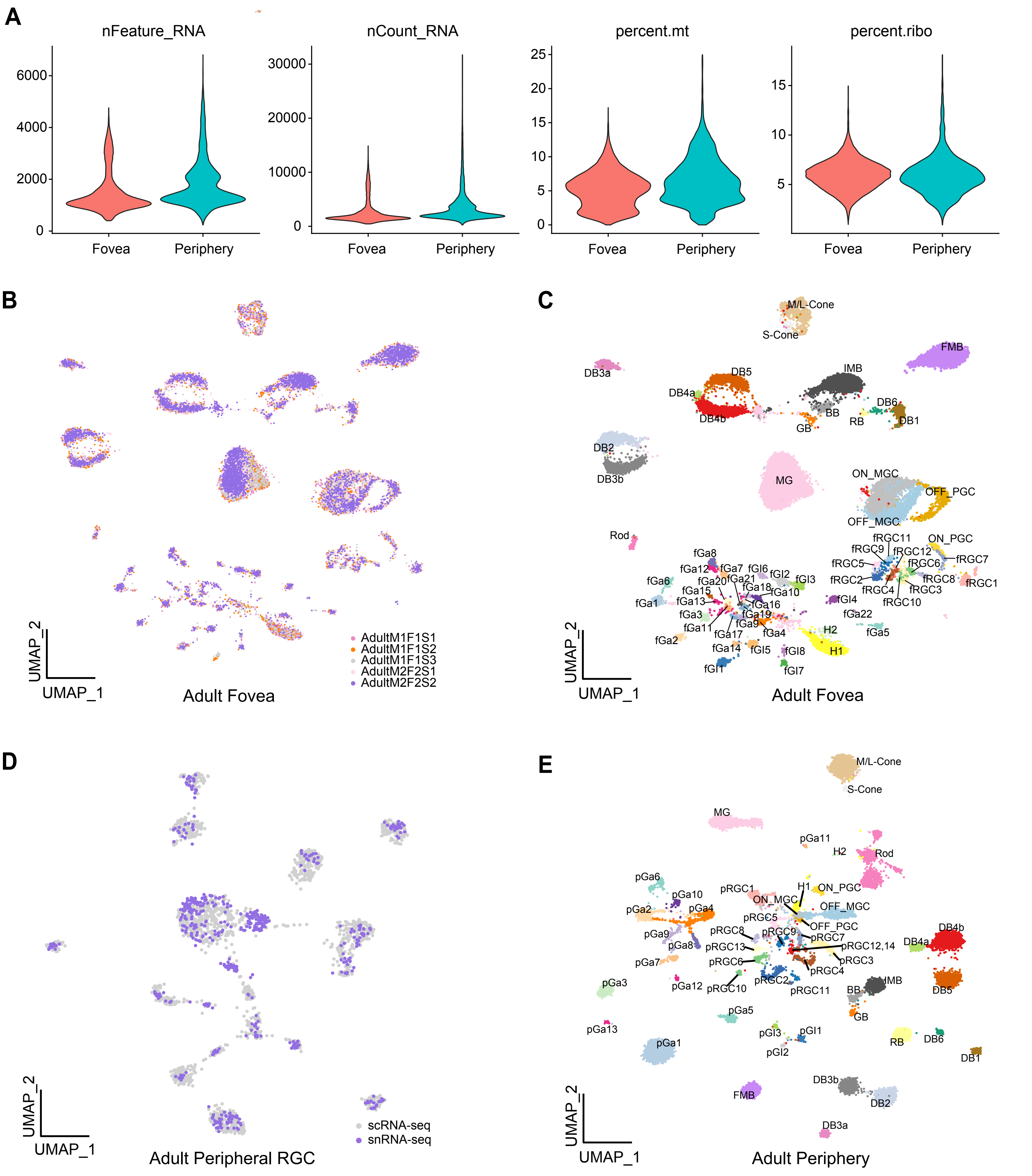

A
nFeature_RNA
0
2000
4000
6000
Fovea
Periphery
nCount_RNA
0
10000
20000
30000
Fovea
Periphery
percent.mt
0
5
10
15
20
25
Fovea
Periphery
percent.ribo
5
10
15
Fovea
Periphery
B
AdultM1F1S1
AdultM1F1S2
AdultM1F1S3
AdultM2F2S1
AdultM2F2S2
UMAP_2
UMAP_1
Adult Fovea
C
M/L-Cone
S-Cone
FMB
DB3a
DB5
IMB
DB4a
DB4b
BB
DB6
RB
DB1
GB
DB2
DB3b
MG
ON_MGC
OFF_PGC
OFF_MGC
Rod
fRGC11
ON_PGC
fRGC9
fRGC12
fRGC7
fRGC5
fRGC6
fRGC2
fRGC8
fRGC1
fRGC4
fRGC3
fRGC10
fGa8
fGa12
fGa7
fGl6
fGl2
fGl3
fGa6
fGa20
fGa21
fGa15
fGa18
fGa10
fGa1
fGa13
fGa16
fGl4
fGa3
fGa19
fGa22
fGa11
fGa9
fGa4
fGa5
fGa17
H2
fGa2
fGa14
fGl5
fGl8
H1
fGl1
fGl7
UMAP_2
UMAP_1
Adult Fovea
D
scRNA-seq
snRNA-seq
UMAP_2
UMAP_1
Adult Peripheral RGC
E
M/L-Cone
S-Cone
MG
pGa11
H2
Rod
pGa6
pGa10
pRGC1
H1
ON_PGC
ON_MGC
pGa2
pGa4
OFF_MGC
pRGC5
OFF_PGC
pGa9
pRGC8
pRGC9
pRGC7
DB4b
pGa8
pRGC13
DB4a
pRGC12,14
pGa7
pRGC6
pRGC3
pRGC4
pGa3
pGa12
pRGC10
pRGC2
IMB
pRGC11
BB
DB5
GB
pGa5
pGa13
pGl3
pGl1
RB
DB6
pGa1
pGl2
DB1
DB3b
FMB
DB2
DB3a
UMAP_2
UMAP_1
Adult Periphery

### Figure S2

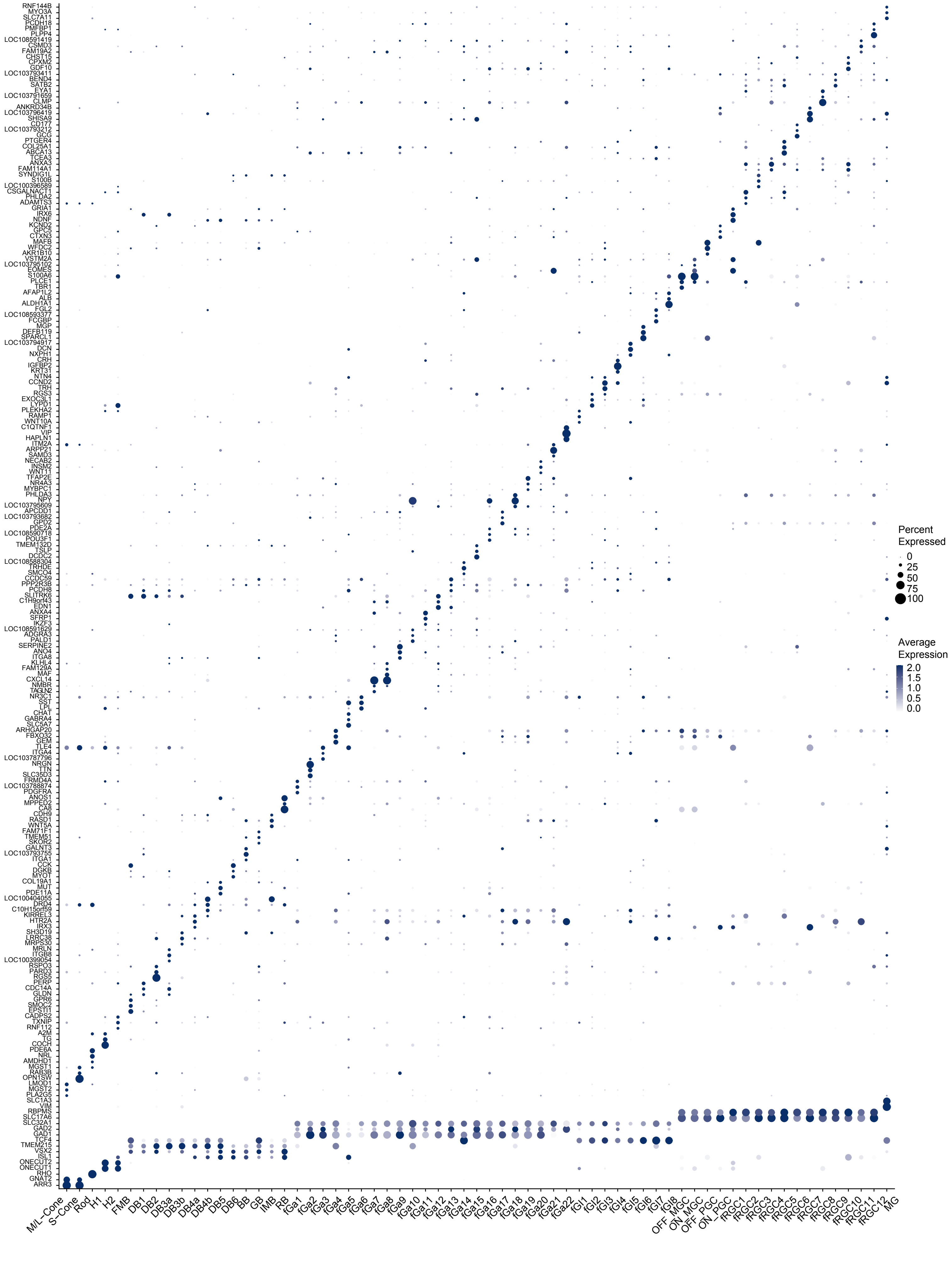
RNF144B
MYO3A
SLC7A11
PCDH18
PMFBP1
PLPP4
LOC108591419
CSMD3
FAM19A2
CHST15
CPXM2
GDF10
LOC103793411
BEND4
SATB2
EYA1
LOC103791659
CLMP
ANKRD34B
LOC103796419
SHISA9
CD177
LOC103793212
GCG
PTGER4
COL25A1
ABCA13
TCEA3
ANXA3
FAM114A1
SYNDIG1L
S100B
LOC100396589
CSGALNACT1
PHLDA2
ADAMTS3
GRIA1
IRX6
NDNF
KCND2
GPC5
CTXN3
MAFB
WFDC2
AKR1B10
VSTM2A
LOC103795102
EOMES
S100A6
PLCE1
TBR1
AFAP1L2
ALB
ALDH1A1
FGL2
LOC108593377
FCGBP
MGP
DEFB119
SPARCL1
LOC103794917
DCN
NXPH1
CRH
IGFBP2
KRT31
NTN4
CCND2
TRH
RGS3
EXOC3L1
LYPD1
PLEKHA2
RAMP1
WNT10A
C1QTNF1
VIP
HAPLN1
ITM2A
ARPP21
SAMD3
NECAB2
INSM2
WNT11
TFAP2E
NR4A3
MYBPC1
PHLDA3
NPY
LOC103795609
APCDD1
LOC103793682
GPD2
PDE2A
LOC108590718
POU3F1
TMEM132D
TSLP
DCDC2
LOC108588304
TRHDE
SMOC4
CCDC59
PPP2R3B
PCDH8
SLITRK6
C1H9orf43
EDN1
ANXA4
SFRP1
IKZF3
LOC108591629
ADGRA3
PALD1
SERPINE2
ANO4
ITGA8
KLHL4
FAM129A
MAF
CXCL14
NMBR
TAGLN2
NR3C1
SST
LPL
CHAT
GABRA4
SLC5A7
ARHGAP20
FBXO32
GEM
TLE4
ITGA4
LOC103787796
NRGN
TTN
SLC35D3
FRMD4A
LOC103788874
PDGFRA
ANOS1
MPPED2
CA8
CDH9
RASD1
WNT5A
FAM71F1
TMEM51
SKOR2
GALNT3
LOC103793755
ITGA1
CCK
DGKB
MYOT
COL19A1
MUT
PDE11A
LOC100404055
DRD4
C10H15orf59
KIRREL3
HTR2A
IRX3
SH3D19
LRRC38
MRPS30
MRLN
ITGB8
LOC100399054
RSPO3
PARD3
RGS5
PERP
CDC14A
GLDN
GPR6
SMOC2
EPSTI1
CADPS2
TXNIP
RNF112
A2M
TG
COCH
PDE6A
NRL
AMDHD1
MGST1
RAB3B
OPN1SW
LMOD1
MGST2
PLA2G5
SLC1A3
VIM
RBPMS
SLC17A6
SLC32A1
GAD2
GAD1
TCF4
TMEM215
VSX2
ISL1
ONECUT2
ONECUT1
RHO
GNAT2
ARR3
M/L-Cone
S-Cone
Rod
H1
H2
FMB
DB1
DB2
DB3a
DB3b
DB4a
DB4b
DB5
DB6
BB
GB
IMB
RB
fGa1
fGa2
fGa3
fGa4
fGa5
fGa6
fGa7
fGa8
fGa9
fGa10
fGa11
fGa12
fGa13
fGa14
fGa15
fGa16
fGa17
fGa18
fGa19
fGa20
fGa21
fGa22
fGl1
fGl2
fGl3
fGl4
fGl5
fGl6
fGl7
fGl8
OFF_MGC
ON_MGC
OFF_PGC
ON_PGC
fRGC1
fRGC2
fRGC3
fRGC4
fRGC5
fRGC6
fRGC7
fRGC8
fRGC9
fRGC10
fRGC11
fRGC12
MG
Percent Expressed
0
25
50
75
100
Average Expression
2.0
1.5
1.0
0.5
0.0

### Figure S3

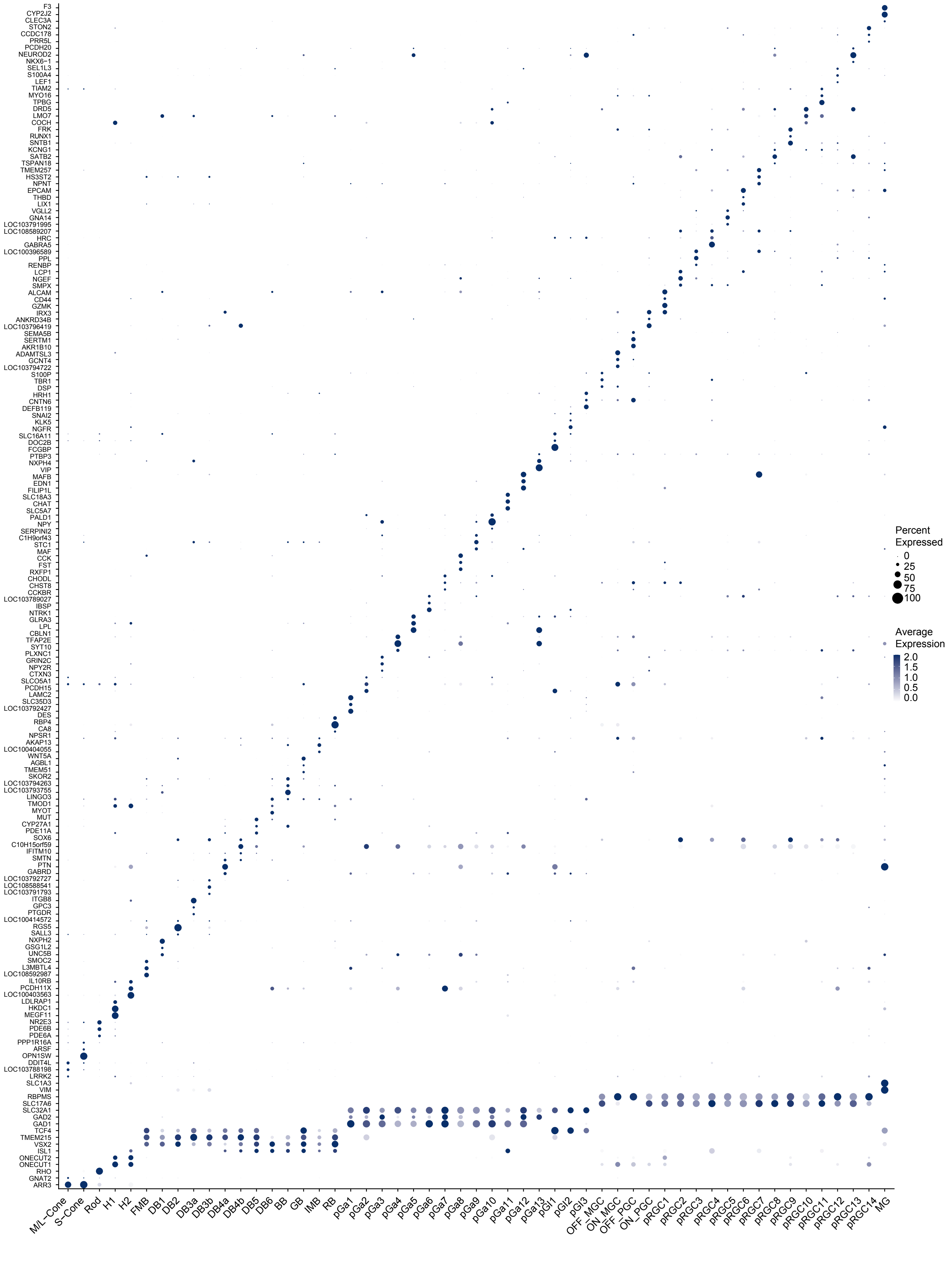
F3
CYP2J2
CLEC3A
STON2
CCDC178
PRR5L
PCDH20
NEUROD2
NKX6−1
SEL1L3
S100A4
LEF1
TIAM2
MYO16
TPBG
DRD5
LMO7
COCH
FRK
RUNX1
SNTB1
KCNG1
SATB2
TSPAN18
TMEM257
HS3ST2
NPNT
EPCAM
THBD
LIX1
VGLL2
GNA14
LOC103791995
LOC108589207
HRC
GABRA5
LOC100396589
PPL
RENBP
LCP1
NGEF
SMPX
ALCAM
CD44
GZMK
IRX3
ANKRD34B
LOC103796419
SEMA5B
SERTM1
AKR1B10
ADAMTSL3
GCNT4
LOC103794722
S100P
TBR1
DSP
HRH1
CNTN6
DEFB119
SNAI2
KLK5
NGFR
SLC16A11
DOC2B
FCGBP
PTBP3
NXPH4
VIP
MAFB
EDN1
FILIP1L
SLC18A3
CHAT
SLC5A7
PALD1
NPY
SERPINI2
C1H9orf43
STC1
MAF
CCK
FST
RXFP1
CHODL
CHST8
CCKBR
LOC103789027
IBSP
NTRK1
GLRA3
LPL
CBLN1
TFAP2E
SYT10
PLXNC1
GRIN2C
NPY2R
CTXN3
SLCO5A1
PCDH15
LAMC2
SLC35D3
LOC103792427
DES
RBP4
CA8
NPSR1
AKAP13
LOC100404055
WNT5A
AGBL1
TMEM51
SKOR2
LOC103794263
LOC103793755
LINGO3
TMOD1
MYOT
MUT
CYP27A1
PDE11A
SOX6
C10H15orf59
IFITM10
SMTN
PTN
GABRD
LOC103792727
LOC108588541
LOC103791793
ITGB8
GPC3
PTGDR
LOC100414572
RGS5
SALL3
NXPH2
GSG1L2
UNC5B
SMOC2
L3MBTL4
LOC108592987
IL10RB
PCDH11X
LOC100403563
LDLRAP1
HKDC1
MEGF11
NR2E3
PDE6B
PDE6A
PPP1R16A
ARSF
OPN1SW
DDIT4L
LOC103788198
LRRK2
SLC1A3
VIM
RBPMS
SLC17A6
SLC32A1
GAD2
GAD1
TCF4
TMEM215
VSX2
ISL1
ONECUT2
ONECUT1
RHO
GNAT2
ARR3
M/L-Cone
S-Cone
Rod
H1
H2
FMB
DB1
DB2
DB3a
DB3b
DB4a
DB4b
DB5
DB6
BB
GB
IMB
RB
pGa1
pGa2
pGa3
pGa4
pGa5
pGa6
pGa7
pGa8
pGa9
pGa10
pGa11
pGa12
pGa13
pGl1
pGl2
pGl3
OFF_MGC
ON_MGC
OFF_PGC
ON_PGC
pRGC1
pRGC2
pRGC3
pRGC4
pRGC5
pRGC6
pRGC7
pRGC8
pRGC9
pRGC10
pRGC11
pRGC12
pRGC13
pRGC14
MG
Percent Expressed
0
25
50
75
100
Average Expression
2.0
1.5
1.0
0.5
0.0

Figure S4

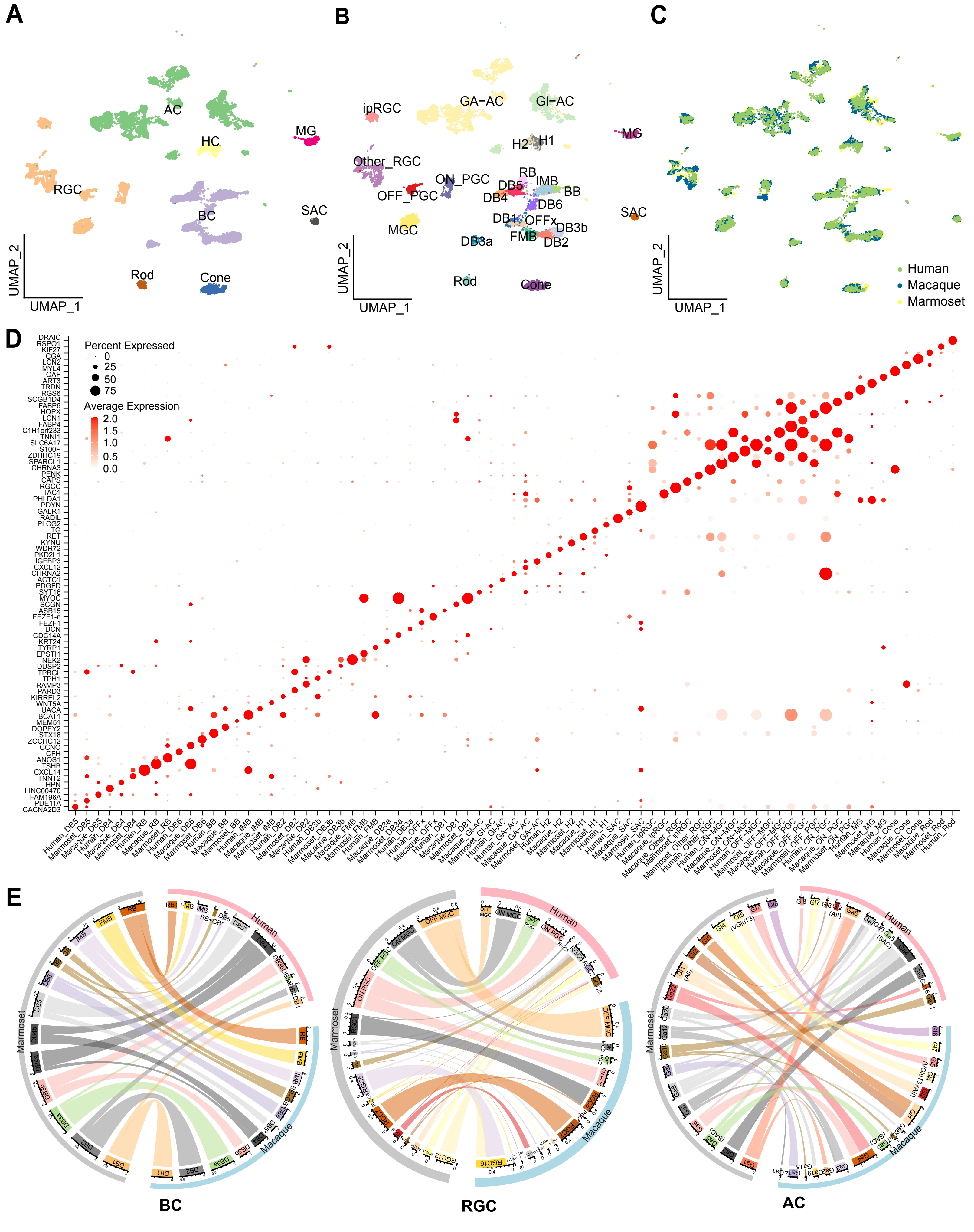
A
AC
HC
MG
RGC
BC
SAC
Rod
Cone
UMAP_2
UMAP_1
B
ipRGC
GA−AC
GI−AC
MG
H2
H1
Other_RGC
RB
ON_PGC
DB5
IMB
OFF_PGC
DB4
BB
DB6
DB1
OFFx
SAC
MGC
DB3b
FMB
DB3a
DB2
Rod
Cone
C
Human
Macaque
Marmoset
D
Percent Expressed
0
25
50
75
Average Expression
2.0
1.5
1.0
0.5
0.0
DRAIC
RSPO1
KIF27
CGA
LCN2
MYL4
OAF
ART3
TRDN
RGS6
SCGB1D4
FABP6
HOPX
LCN1
FABP4
C1H1orf233
TNNI1
SLC6A17
S100P
ZDHHC19
SPARCL1
CHRNA3
PENK
CAPS
RGCC
TAC1
PHLDA1
PDYN
GALR1
RADIL
PLCG2
TG
RET
KYNU
WDR72
PKD2L1
IGFBP3
CXCL12
CHRNA2
ACTC1
PDGFD
SYT16
MYOC
SCGN
ASB15
FEZF1-n
FEZF1
DCN
CDC14A
KRT24
TYRP1
EPSTI1
NEK2
DUSP2
TPBGL
TPH1
RAMP3
PARD3
KIRREL2
WNT5A
UACA
BCAT1
TMEM51
DOPEY2
STX18
ZCCHC12
CCNO
CFH
ANOS1
TSHB
CXCL14
TNNT2
HPN
LINC00470
FAM196A
PDE11A
CACNA2D3
E
Human
Marmoset
Macaque
BC
RGC
AC

**Figure S5**

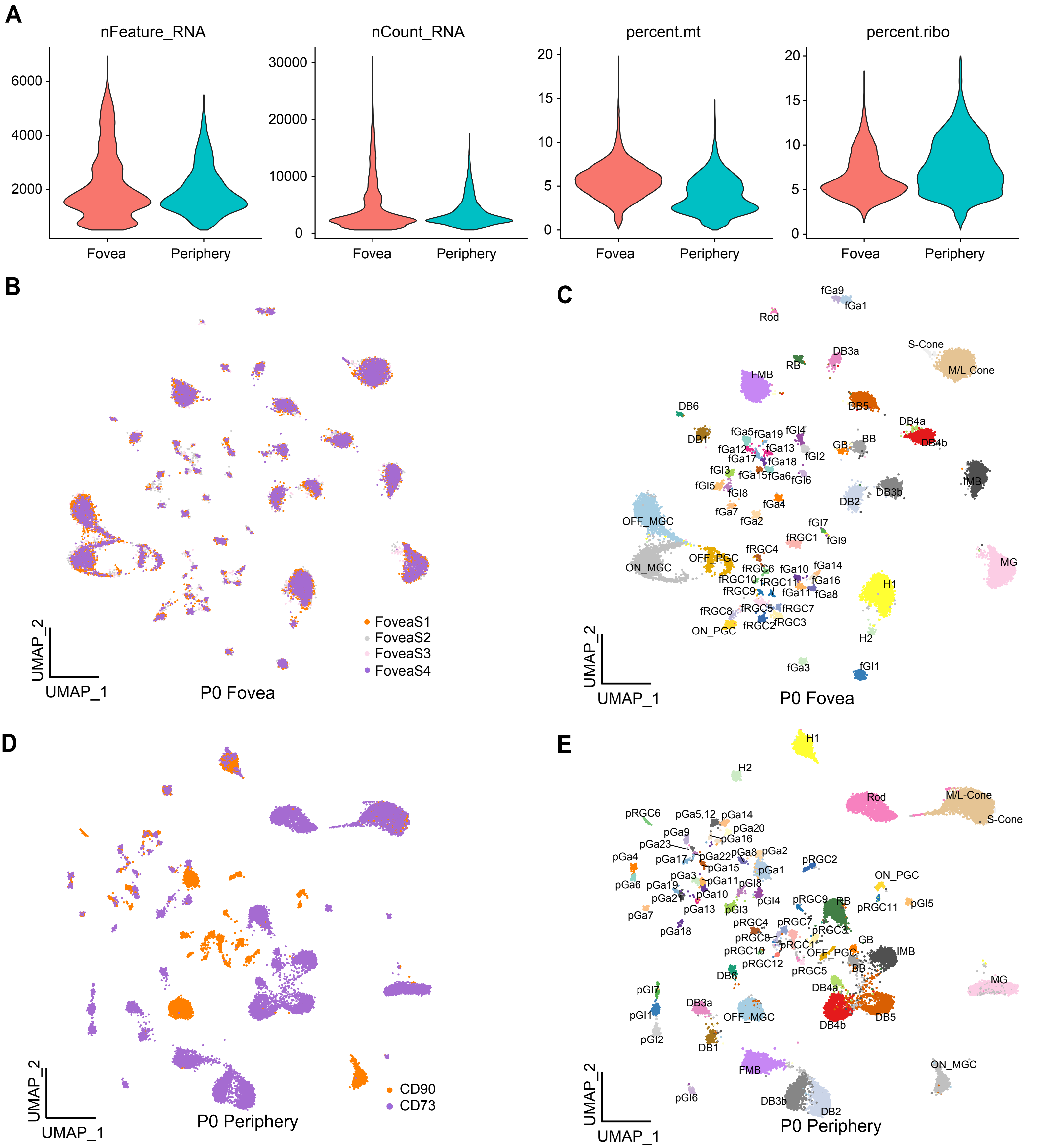
A
nFeature_RNA
nCount_RNA
percent.mt
percent.ribo
6000
4000
2000
30000
20000
10000
0
20
15
10
5
0
Fovea
Periphery
B
FoveaS1
FoveaS2
FoveaS3
FoveaS4
UMAP_2
UMAP_1
P0 Fovea
C
fGa9
fGa1
Rod
S-Cone
M/L-Cone
DB3a
RB
FMB
DB5
DB6
DB4a
DB4b
DB1
fGa5
fGa19
fGl4
fGa12
fGa13
fGa17
fGa18
fGl2
GB
BB
fGl3
fGa15
fGa6
fGl6
IMB
fGl5
fGl8
DB2
DB3b
fGa7
fGa4
fGa2
OFF_MGC
fGl7
fRGC1
fGl9
fRGC4
MG
OFF_PGC
ON_MGC
fRGC6
fGa10
fGa14
fRGC10
fRGC11
fGa16
H1
fRGC9
fGa11
fGa8
fRGC8
fRGC5
fRGC7
fRGC2
fRGC3
ON_PGC
H2
fGa3
fGl1
P0 Fovea
D
CD90
CD73
P0 Periphery
E
H1
H2
Rod
M/L-Cone
S-Cone
pRGC6
pGa5,12
pGa14
pGa9
pGa20
pGa16
pGa23
pGa22
pGa8
pGa2
pGa4
pGa17
pGa15
pGa1
pRGC2
pGa3
pGa6
pGa19
pGa11
pGl8
ON_PGC
pGa21
pGa10
pGl4
pRGC9
RB
pRGC11
pGl5
pGa13
pGl3
pGa7
pRGC4
pRGC7
pGa18
pRGC8
pRGC3
GB
pRGC1
pRGC10
OFF_PGC
IMB
pRGC12
BB
DB6
pRGC5
MG
DB4a
pGl7
DB3a
pGl1
OFF_MGC
DB4b
DB5
pGl2
DB1
FMB
ON_MGC
pGl6
DB3b
DB2
P0 Periphery

### Figure S6

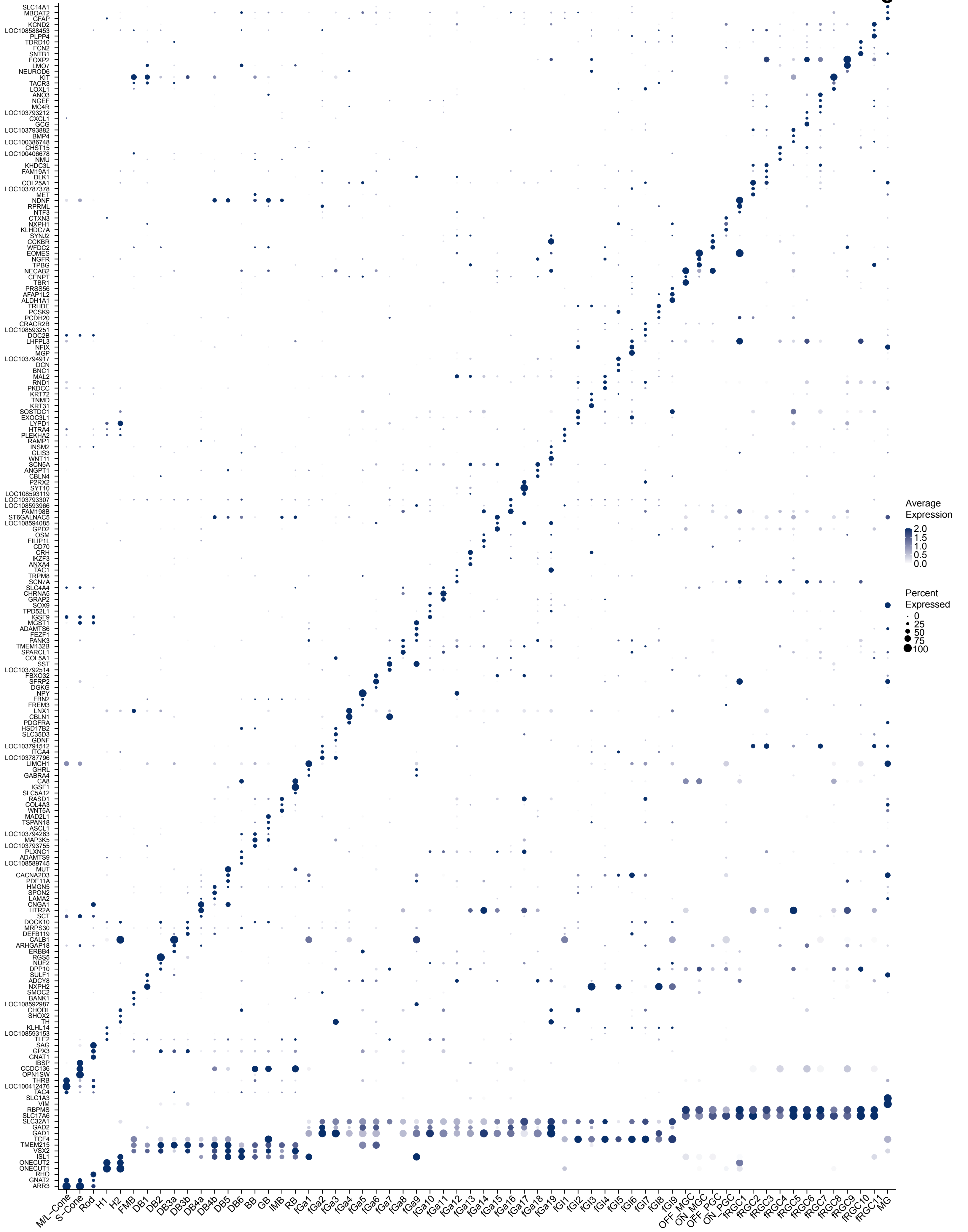
Average Expression
2.0
1.5
1.0
0.5
0.0
Percent Expressed
0
25
50
75
100
M/L-Cone
S-Cone
Rod
H1
H2
FMB
DB1
DB2
DB3a
DB3b
DB4a
DB4b
DB5
DB6
BB
GB
IMB
RB
fGa1
fGa2
fGa3
fGa4
fGa5
fGa6
fGa7
fGa8
fGa9
fGa10
fGa11
fGa12
fGa13
fGa14
fGa15
fGa16
fGa17
fGa18
fGa19
fGl1
fGl2
fGl3
fGl4
fGl5
fGl6
fGl7
fGl8
fGl9
OFF_MGC
ON_MGC
OFF_PGC
ON_PGC
fRGC1
fRGC2
fRGC3
fRGC4
fRGC5
fRGC6
fRGC7
fRGC8
fRGC9
fRGC10
fRGC11
MG

### Figure S7

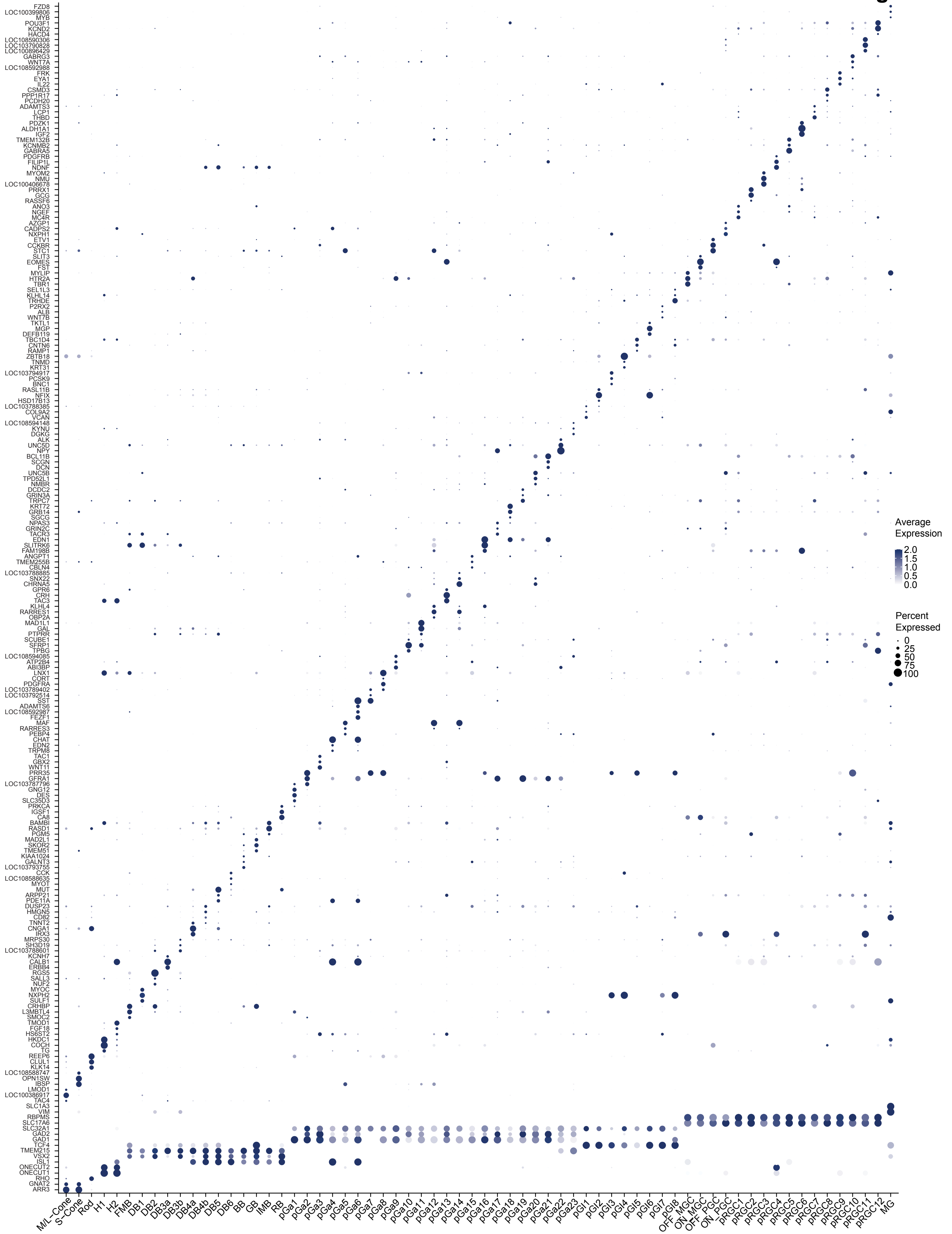
FZD8
LOC100399806
MYB
POU3F1
KCND2
HACD4
LOC108590306
LOC103790828
LOC100896429
GABRG3
WNT7A
LOC108592988
FRK
EYA1
IL22
CSMD3
PPP1R17
PCDH20
ADAMTS3
LCP1
THBD
PDZK1
ALDH1A1
IGF2
TMEM132B
KCNMB2
GABRA5
PDGFRB
FILIP1L
NDNF
MYOM2
NMU
LOC100406678
PRRX1
GCG
RASSF6
ANO3
NGEF
MC4R
AZGP1
CADPS2
NXPH1
ETV1
CCKBR
STC1
SLIT3
EOMES
FST
MYLIP
HTR2A
TBR1
SEL1L3
KLHL14
TRHDE
P2RX2
ALB
WNT7B
TKTL1
MGP
DEFB119
TBC1D4
CNTN6
RAMP1
ZBTB18
TNMD
KRT31
LOC103794917
PCSK9
BNC1
RASL11B
NFIX
HSD17B13
LOC103788385
COL9A2
VCAN
LOC108594148
KYNU
DGKG
ALK
UNC5D
NPY
BCL11B
SCGN
DCN
UNC5B
TPD52L1
NMBR
DCDC2
GRIN3A
TRPC7
KRT72
GRB14
SGCG
NPAS3
GRIN2C
TACR3
EDN1
SLITRK6
FAM198B
ANGPT1
TMEM255B
CBLN4
LOC103788885
SNX22
CHRNA5
GPR6
CRH
TAC3
KLHL4
RARRES1
OBP2A
MAD1L1
GAL
PTPRR
SCUBE1
SFRP1
TPBG
LOC108594085
ATP2B4
ABI3BP
LNX1
CORT
PDGFRA
LOC103789402
LOC103792514
SST
ADAMTS6
LOC108592987
FEZF1
MAF
RARRES3
PEBP4
CHAT
EDN2
TRPM8
TAC1
GBX2
WNT11
PRR35
GFRA1
LOC103787796
GNG12
DES
SLC35D3
PRKCA
IGSF1
CA8
BAMBI
RASD1
PGM5
MAD2L1
SKOR2
TMEM51
KIAA1024
GALNT3
LOC103793755
CCK
LOC108588635
MYOT
MUT
ARPP21
PDE11A
DUSP23
HMGN5
CD82
TNNT2
CNGA1
IRX3
MRPS30
SH3D19
LOC103788601
KCNH7
CALB1
ERBB4
RGS5
SALL3
NUF2
MYOC
NXPH2
SULF1
CRHBP
L3MBTL4
SMOC2
TMOD1
HS6ST2
FGF18
HKDC1
COCH
TG
REEP6
CLUL1
KLK14
LOC108588747
OPN1SW
IBSP
LMOD1
LOC100386917
TAC4
SLC1A3
VIM
RBPMS
SLC17A6
SLC32A1
GAD2
GAD1
TCF4
TMEM215
VSX2
ISL1
ONECUT2
ONECUT1
RHO
GNAT2
ARR3
M/L-Cone
S-Cone
Rod
H1
H2
FMB
DB1
DB2
DB3a
DB3b
DB4a
DB4b
DB5
DB6
BB
GB
IMB
RB
pGa1
pGa2
pGa3
pGa4
pGa5
pGa6
pGa7
pGa8
pGa9
pGa10
pGa11
pGa12
pGa13
pGa14
pGa15
pGa16
pGa17
pGa18
pGa19
pGa20
pGa21
pGa22
pGa23
pGl1
pGl2
pGl3
pGl4
pGl5
pGl6
pGl7
pGl8
OFF MGC
ON MGC
OFF PGC
ON PGC
pRGC1
pRGC2
pRGC3
pRGC4
pRGC5
pRGC6
pRGC7
pRGC8
pRGC9
pRGC10
pRGC11
pRGC12
MG
Average Expression
2.0
1.5
1.0
0.5
0.0
Percent Expressed
0
25
50
75
100

**Figure S8**

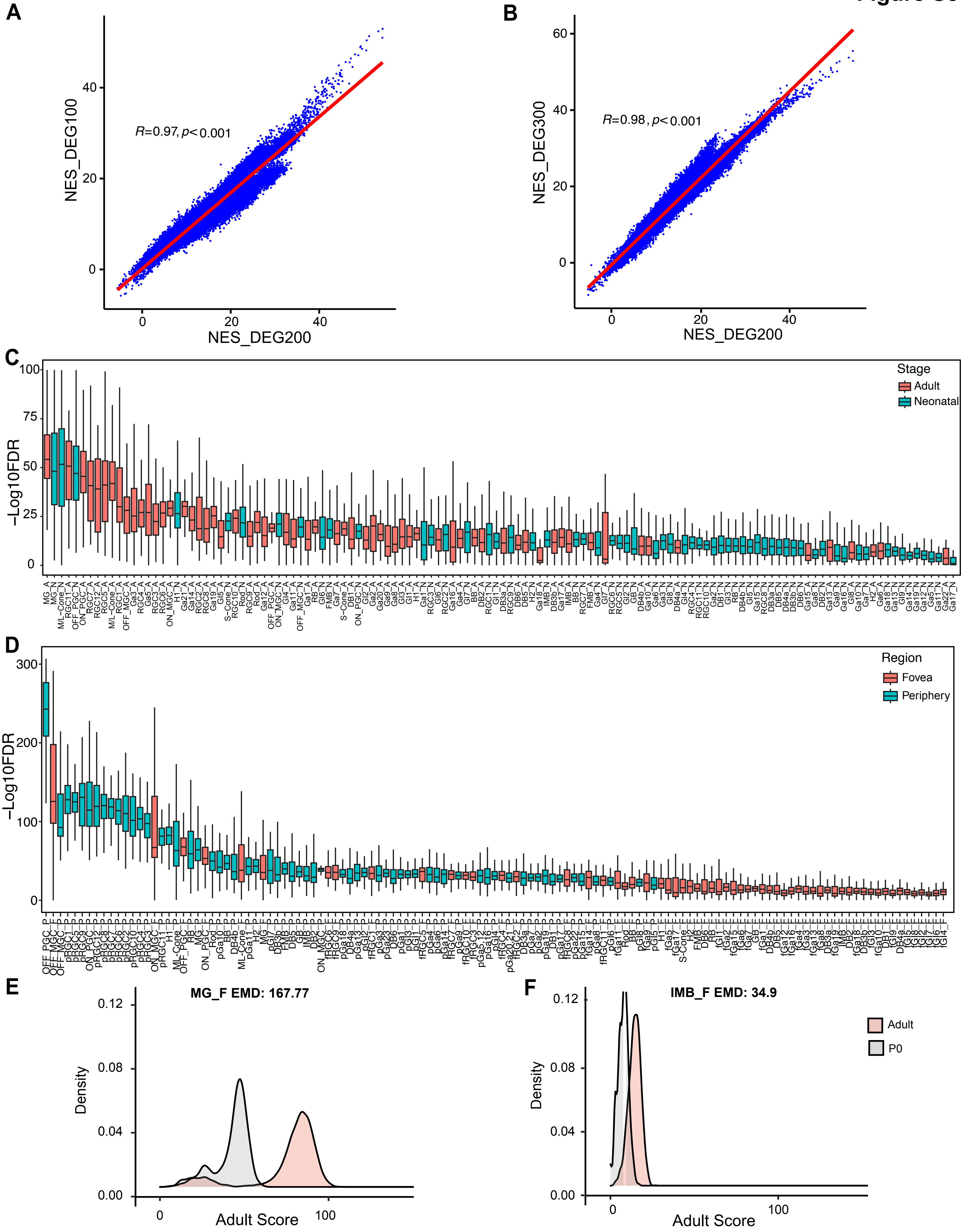
A
B
C
D
E
F
R=0.97, p< 0.001
R=0.98, p< 0.001
NES_DEG100
NES_DEG300
NES_DEG200
-Log10FDR
Stage
Adult
Neonatal
Region
Fovea
Periphery
MG_F EMD: 167.77
IMB_F EMD: 34.9
Density
Adult Score
Adult
P0

### Figure S9

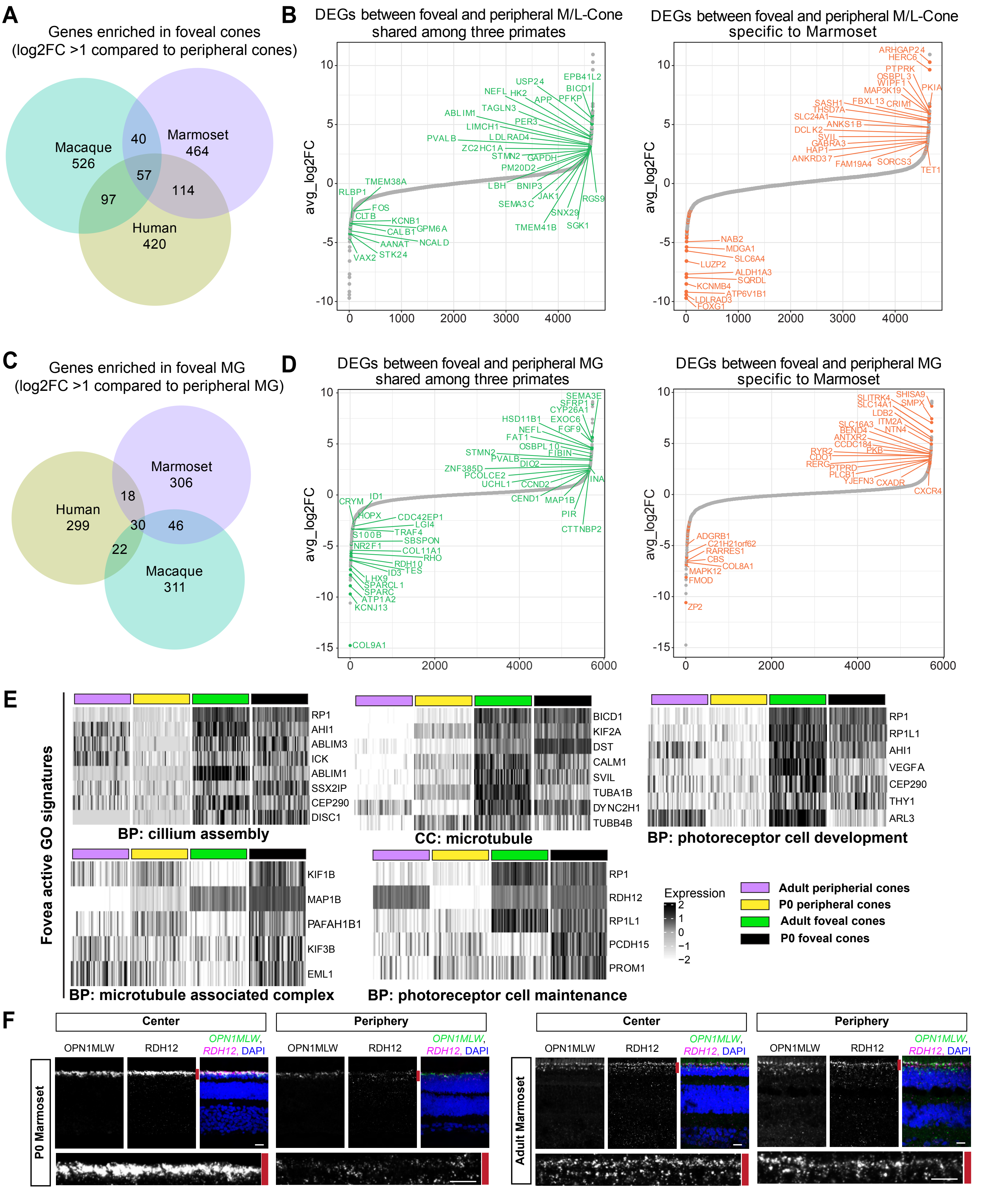
A
Genes enriched in foveal cones
(log2FC >1 compared to peripheral cones)
Macaque 526
Marmoset 464
Human 420
40
57
97
114
B
DEGs between foveal and peripheral M/L-Cone shared among three primates
DEGs between foveal and peripheral M/L-Cone specific to Marmoset
avg_log2FC
C
Genes enriched in foveal MG
(log2FC >1 compared to peripheral MG)
Marmoset 306
Human 299
Macaque 311
18
30
46
22
D
DEGs between foveal and peripheral MG shared among three primates
DEGs between foveal and peripheral MG specific to Marmoset
E
Fovea active GO signatures
BP: cillium assembly
CC: microtubule
BP: photoreceptor cell development
BP: microtubule associated complex
BP: photoreceptor cell maintenance
Expression
Adult peripherial cones
P0 peripheral cones
Adult foveal cones
P0 foveal cones
F
Center
Periphery
OPN1MLW
RDH12
OPN1MLW, RDH12, DAPI
P0 Marmoset
Adult Marmoset

### Figure S10

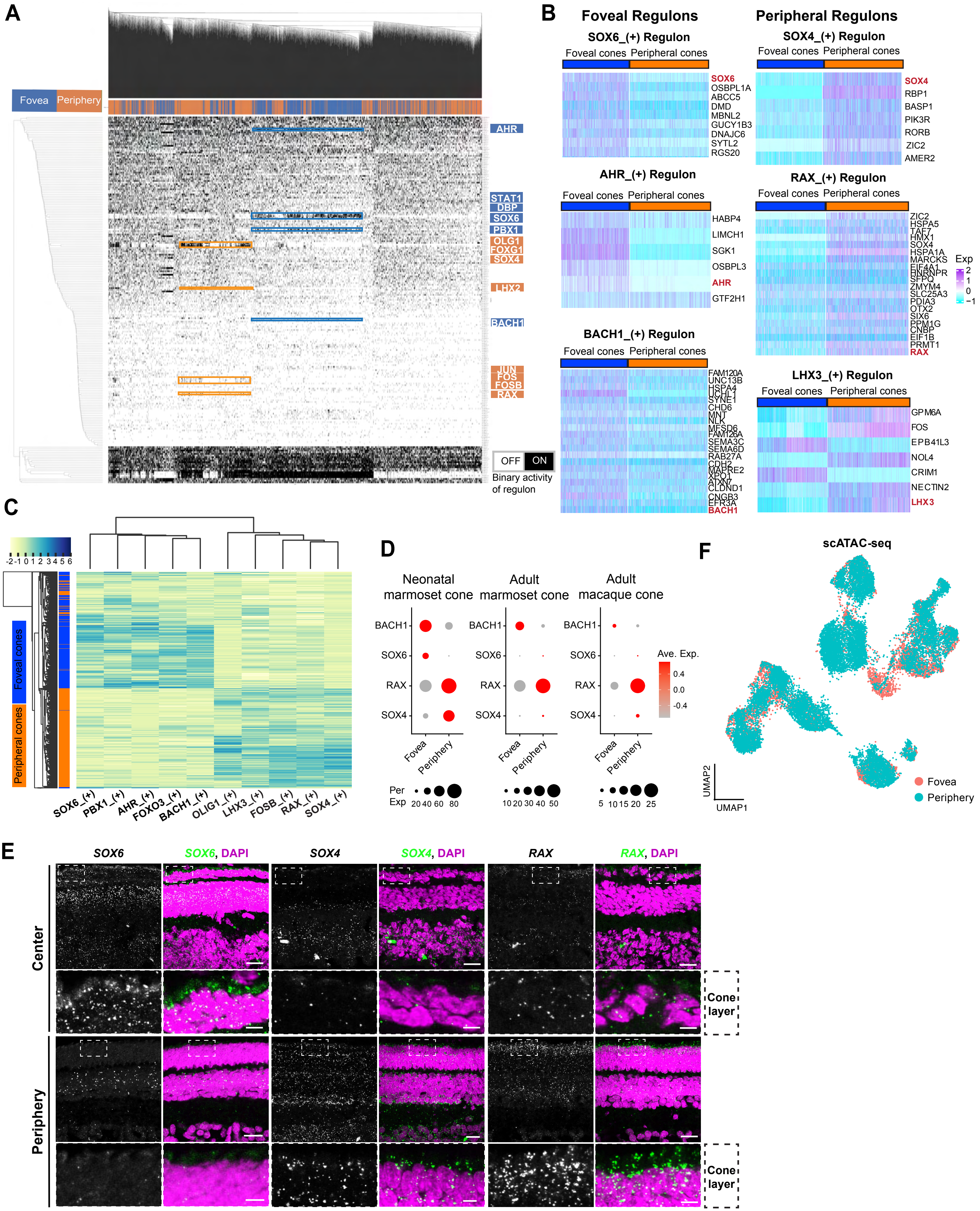
A
Fovea
Periphery
AHR
STAT1
DBP
SOX6
PBX1
OLG1
FOXG1
SOX4
LHX2
BACH1
JUN
FOS
FOSB
RAX
OFF
ON
Binary activity of regulon
B
Foveal Regulons
Peripheral Regulons
SOX6_(+) Regulon
Foveal cones
Peripheral cones
SOX6
OSBPL1A
ABCC5
DMD
MBNL2
GUCY1B3
DNAJC6
SYTL2
RGS20
SOX4_(+) Regulon
SOX4
RBP1
BASP1
PIK3R
RORB
ZIC2
AMER2
AHR_(+) Regulon
HABP4
LIMCH1
SGK1
OSBPL3
AHR
GTF2H1
RAX_(+) Regulon
Exp
BACH1_(+) Regulon
LHX3_(+) Regulon
GPM6A
FOS
EPB41L3
NOL4
CRIM1
NECTIN2
LHX3
C
Foveal cones
Peripheral cones
SOX6_(+)
PBX1_(+)
AHR_(+)
FOXO3_(+)
BACH1_(+)
OLIG1_(+)
LHX3_(+)
FOSB_(+)
RAX_(+)
SOX4_(+)
D
Neonatal marmoset cone
Adult marmoset cone
Adult macaque cone
BACH1
SOX6
RAX
SOX4
Fovea
Periphery
Ave. Exp.
Per Exp
F
scATAC-seq
UMAP1
UMAP2
Fovea
Periphery
E
SOX6
SOX6, DAPI
SOX4
SOX4, DAPI
RAX
RAX, DAPI
Center
Periphery
Cone layer

### Figure S11

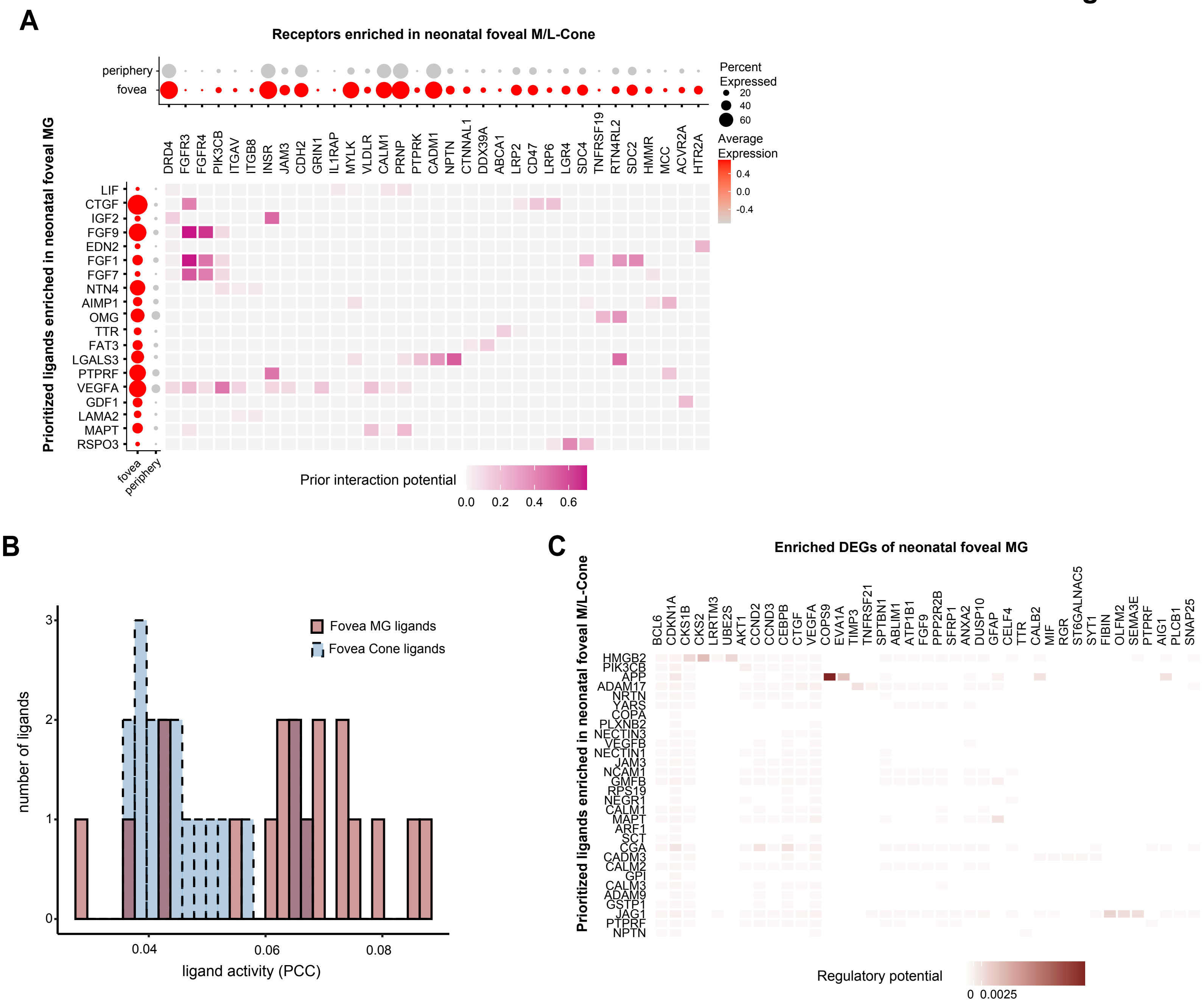
A
Receptors enriched in neonatal foveal M/L-Cone
periphery
fovea
Percent Expressed
20
40
60
Average Expression
0.4
0.0
-0.4
DRD4 FGFR3 FGFR4 PIK3CB ITGAV ITGB8 INSR JAM3 CDH2 GRIN1 IL1RAP MYLK VLDLR CALM1 PRNP PTPRK CADM1 NPTN CTNNAL1 DDX39A ABCA1 LRP2 CD47 LRP6 LGR4 SDC4 TNFRSF19 RTN4RL2 SDC2 HMMR MCC ACVR2A HTR2A
Prioritized ligands enriched in neonatal foveal MG
LIF CTGF IGF2 FGF9 EDN2 FGF1 FGF7 NTN4 AIMP1 OMG TTR FAT3 LGALS3 PTPRF VEGFA GDF1 LAMA2 MAPT RSPO3
fovea
periphery
Prior interaction potential
0.0
0.2
0.4
0.6
B
number of ligands
ligand activity (PCC)
Fovea MG ligands
Fovea Cone ligands
0
1
2
3
0.04
0.06
0.08
C
Enriched DEGs of neonatal foveal MG
Prioritized ligands enriched in neonatal foveal M/L-Cone
BCL6 CDKN1A CKS1B CKS2 LRRTM3 UBE2S AKT1 CCND2 CCND3 CEBPB CTGF VEGFA COPS9 EVA1A TIMP3 TNFRSF21 SPTBN1 ABLIM1 ATP1B1 FGF9 PPP2R2B SFRP1 ANXA2 DUSP10 GFAP CELF4 TTR CALB2 MIF RGR ST6GALNAC5 SYT1 FIBIN OLFM2 SEMA3E PTPRF AIG1 PLCB1 SNAP25
HMGB2 PIK3CB APP ADAM17 NRTN YARS COPA PLXNB2 NECTIN3 VEGFB NECTIN1 JAM3 NCAM1 GMFB RPS19 NEGR1 CALM1 MAPT ARF1 SCT CGA CADM3 CALM2 GPI CALM3 ADAM9 GSTP1 JAG1 PTPRF NPTN
Regulatory potential
0
0.0025
